# BEguided: Systematic guide and base editor evaluation for precision single-nucleotide editing

**DOI:** 10.64898/2026.11.20.767841

**Authors:** Lukas Theo Schmitt, Pascal Wang, Sara Orugo Medina, Moustafa Abohawya, Frank Buchholz, Shady Sayed

**Affiliations:** Medical Systems Biology, Faculty of Medicine Carl Gustav Carus, TU Dresden, Dresden, Germany.; German Cancer Consortium (DKTK), partner site Dresden and German Cancer Research Center (DKFZ), Heidelberg, Germany.; OncoRay – National Center for Radiation Research in Oncology, Faculty of Medicine and University Hospital Carl Gustav Carus, Technische Universität Dresden and Helmholtz-Zentrum Dresden-Rossendorf, Dresden, Germany.; National Center for Tumor Diseases (NCT), NCT/UCC Dresden, a partnership between DKFZ, Faculty of Medicine and University Hospital Carl Gustav Carus, TUD Dresden University of Technology, and Helmholtz-Zentrum Dresden-Rossendorf (HZDR), Dresden, Germany.; Mildred Scheel Early Career Center (MSNZ P2), Dresden, Germany. Faculty of Medicine Carl Gustav Carus, TU Dresden, Dresden, Germany.

## Abstract

Base editing enables precise nucleotide substitutions without introducing DNA double-strand breaks, but identifying optimal editor–guide combinations remains challenging because PAM compatibility, editing-window positioning, bystander modifications, and off-target effects must be considered simultaneously. Here, we introduce BEguided, an integrated computational platform for variant-specific guide RNA design and systematic comparison across 38 predefined base-editor profiles, with support for user-defined editors. BEguided incorporates single-nucleotide variants directly into target sequences, eliminating the need for customized reference genomes, while its companion tool, BEguidedOFF, provides base-editing-aware off-target prediction. To demonstrate the utility of this approach, we systematically assessed the accessibility of the 1,000 most frequent ABE-editable cancer mutations to adenine base editing. Under stringent guide-quality, bystander-editing, and predicted off-target criteria, only 7 mutations could be targeted using SpCas9-ABE8e. Expanding the analysis to additional SpCas9-compatible adenine base editors increased this number to 30, while inclusion of SaCas9, relaxed-PAM Cas9-NG/SpG variants, and PAM-less SpRY further expanded the number of potentially targetable mutations to 100, 156, and 238, respectively. These findings highlight the substantial impact of editor selection on mutation accessibility and establish BEguided as a versatile framework for identifying and prioritizing base-editing strategies for disease-associated variants. BEguided is available at https://beguided.buchholzlab.org/.

Figure 1:
Graphical abstract

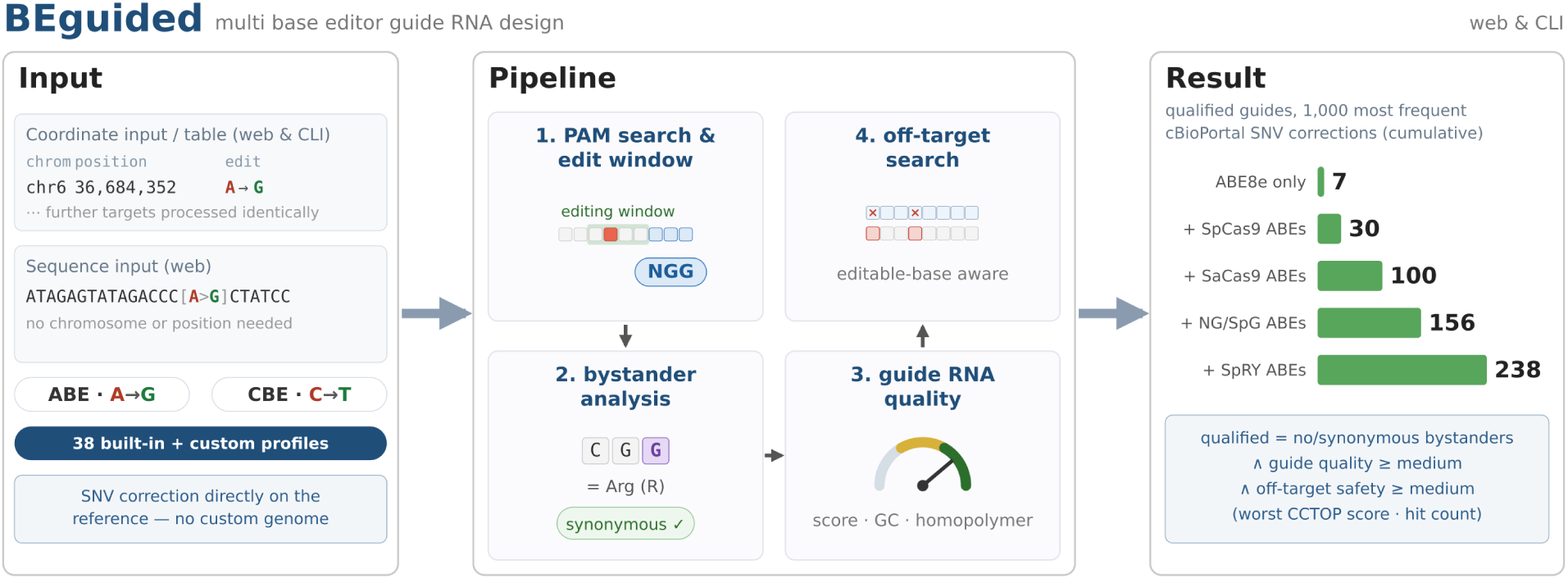

## Introduction

CRISPR base editors enable programmable single-nucleotide conversions without inducing double-strand DNA breaks (1–3). By fusing a catalytically impaired Cas9 nickase to a deaminase enzyme, cytosine base editors (CBEs) and adenine base editors (ABEs) mediate C→T and A→G transitions, respectively. This enables us to model or correct a substantial fraction of pathogenic point mutations. Gaudelli et al. determined that approximately 48% of all pathogenic single-nucleotide variants (SNVs) could be addressed by ABEs (3). Base editing is already delivering on this potential in clinical trials: durable LDL-cholesterol lowering upon *PCSK9* inactivation (4–6), transfusion independence in *β*-thalassemia patients (7), and personalized correction of a patient’s *CPS1* deficiency variant (8). Of particular relevance to this work is the correction of cancer mutations. Many recurrent SNVs are experimentally addressable, as demonstrated for *KRAS* G12D, *TP53* R175H, *TP53* R273H, and *PTEN* R233* in cancer cell lines and PDOs (9, 10). This establishes base editing as a strategy for both functional interrogation and therapeutic mutation correction, a concept recently outlined as a platform for precision oncology (11).

Despite their promise, base editing approaches face several limitations. One of the main challenges lies in the design of optimal guide RNAs (gRNAs) as it is substantially more complex than for conventional nuclease-based Cas9 editing applications. A designer must simultaneously consider whether (i) a suitable PAM is available near the target base; (ii) the target base falls within the editing window of the chosen base editor; (iii) potential bystander editing is identified and assessed for unwanted codon changes; and (iv) the off-target potential is acceptable. These constraints are interdependent: a guide with an otherwise optimal PAM may introduce a nonsynonymous bystander edit, whereas a bystander-free guide may have numerous potential genomic off-targets.

Similarly, an expanding range of base editor variants with distinct properties is becoming available. Variants can vary in PAM sequence (SpCas9-ABE8e with NGG PAM versus SaCas9-ABE8e with NNGRRT PAM), editing window, activity and specificity. Consequently, for optimal experimental design it is necessary to use multiple base editor profiles to determine the best guide RNA and editor combinations.

Existing tools such as BE-Designer (12), ACEofBASEs (13), and BEscreen (14) address these complications only partially. Although they are capable of bulk processing targets, they are restricted to one base editor at a time. Additionally, BE-Designer and ACEofBASEs always provide all potential edits in a given sequence, requiring the user to identify the edits of interest. BEscreen, on the other hand, allows defining specific variants, but it only supports bases that are present in the reference, so designing guides to correct SNVs is not possible. In summary, none of the existing tools provide an integrated workflow for precise, codon-aware, multiplex, multi-editor guide design.

To address these shortcomings, we developed **BEguided** and **BEguidedOFF**, an open-source Python pipeline that unifies guide design, bystander analysis, gRNA evaluation, and off-target scoring in a simple web or command-line interface (CLI). BEguided delegates genome-wide off-target enumeration to BEguidedOFF, a companion library specifically designed for base editing. Together, these tools enable researchers to screen hundreds of targets across multiple editor profiles, producing a unified, comparable table of candidate guides. BEguided can be used as a command-line tool or through its web application, which offers a limited fast off-target search instead of the extensive genome-wide enumeration. We demonstrate the utility of this framework through a large-scale survey of recurrent cancer mutations from the cBioPortal database, quantifying how many mutations allow precise editing when multiple base editors are considered.

## Materials and methods

### Pipeline architecture

BEguided executes in five integrated stages (Figure 2):

1. **Guidefinder.** Given a list of target variants and a reference genome, BEguided identifies all gRNAs that place the target base within the editor’s activity window. It supports all defined base-editor profiles based on a profile file that can be adapted to any base editor (ABE8e, ABE9, CBE6d, …). The profile file specifies two editing windows, a high-activity window, defining the positions at which editing efficiency is the highest, and a broader general activity window, defining positions at which editing may occur at lower efficiency. The high-activity window is used to determine where the on-target editing should happen, while the general activity window is used to determine where bystander editing can happen. To design the guide, for each target the variant base is substituted into the extracted genomic context, enabling rapid single-mutation screening without a modified reference genome. Finally, candidate guides are evaluated using the CRISPRater scoring model, as implemented in CCTop (15).
2. **Bystander analysis.** For every candidate guide, the pipeline lists each bystander position, based on the general edit window defined in the profile file, and simulates editing on these positions. Predicted bystander edits are translated using the user-provided GFF annotation file and its canonical-transcript tag (MANE Select for the human Ensembl annotation, Ensembl canonical for the mouse annotation). Based on the resulting amino acids, it is then determined if the bystander edits would result in a synonymous or nonsynonymous amino acid change. If the target base or a bystander lies outside an annotated coding DNA sequence (CDS), no codon-level consequence can be assigned and the guide is reported as non-coding.
3. **Filtering.** Before proceeding to the computationally expensive off-target search, the guides are filtered according to user-set parameters. There are options for filtering according to bystander editing, guide scores, GC-content, homopolymers, and a trinucleotide edit-context score (expected ABE efficiency of the sequence flanking the edit; see below). This pre-filtering reduces the number of guides subjected to genome-wide off-target analysis and thereby improves computational efficiency.
4. **Off-target search (optional).** BEguided hands the filtered guides to BEguidedOFF (described below), which searches the genome for editable off-target sites and returns a per-guide off-target count and the score of the worst off-target.
5. **Merge & reporting.** Off-target data and the filtered Guidefinder and Bystander analysis results are merged into the main results table.

**Figure 2:**
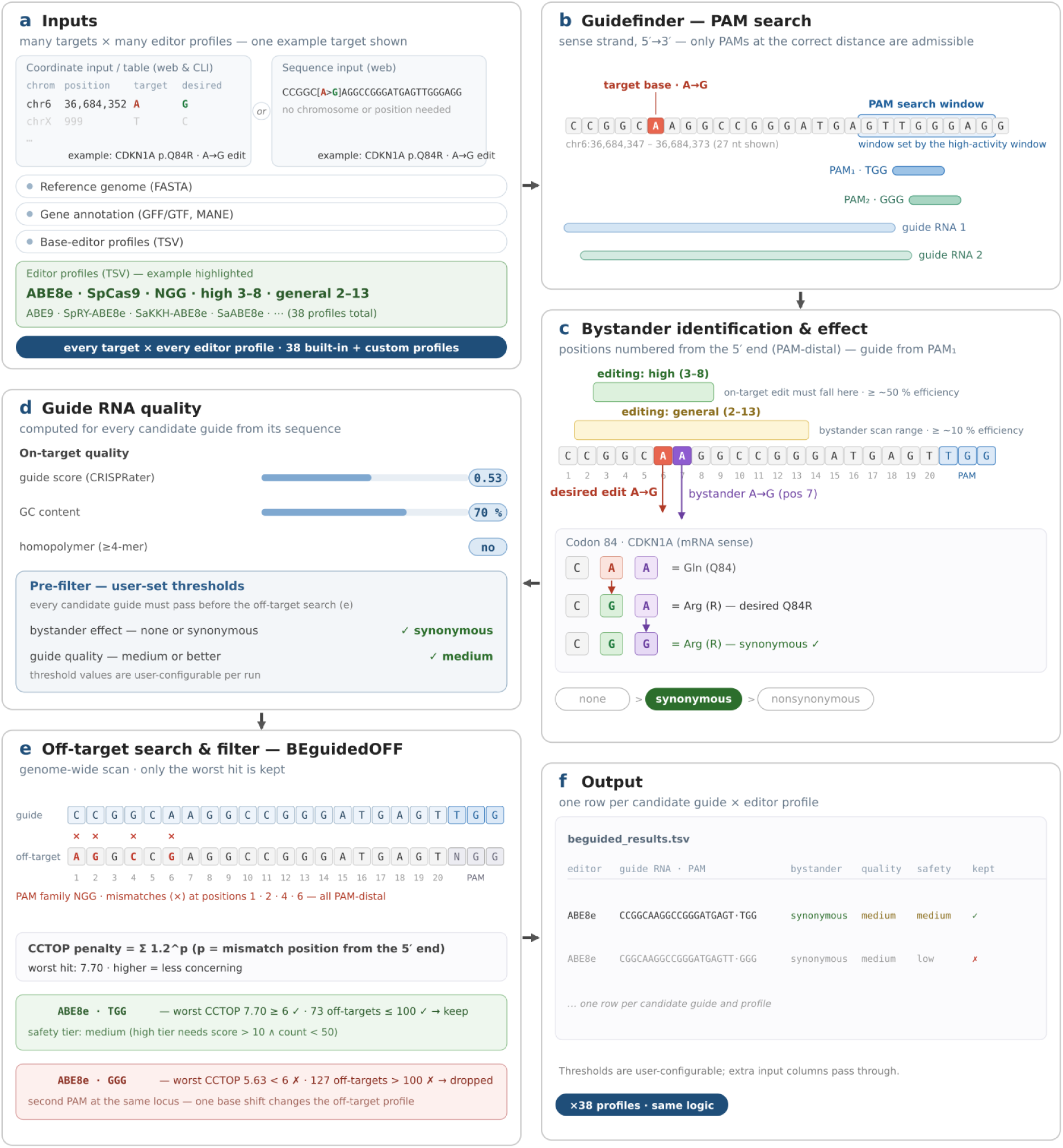
Overview of the BEguided pipeline, illustrated with a real example target. (*CDKN1A* p.Q84R, chr6:36,684,352, A→G restoration edit, ABE8e). **(a)** Inputs: coordinate input / table (web & CLI) or, in the web tool only, a target sequence with the desired edit marked inline; a reference genome (FASTA), a gene annotation (GFF/GTF, MANE), and a base-editor profile table (TSV); every target is processed against every selected profile (38 built-in profiles plus custom profiles). **(b)** Guidefinder: on the sense strand, only PAMs that place the target base within the search window set by the profile’s high-activity window are enumerated and define candidate 20-nt guide RNAs. **(c)** Bystander analysis: guide positions are numbered from the 5′ end; the desired edit and every editable base in the general window are simulated and translated — here the target A→G gives the desired Q84R (CGA), and the bystander A→G at position 7 leaves the codon coding for Arg (CGG, synonymous). **(d)** On-target guide quality (guide score, GC content, homopolymer status) is evaluated, and a pre-filter applying user-set thresholds (bystander effect none or synonymous; guide quality medium or better) removes candidates before the off-target search. **(e)** Off-target search: BEguidedOFF scans the genome on the GPU, keeps only the worst hit per guide and scores it with the CCTop penalty Σ 1.2^p; guides failing the user-set safety thresholds are dropped. **(f)** Output: surviving guides are merged into beguided_results.tsv, one row per candidate guide × editor profile; thresholds are user-configurable and extra input columns pass through to the results.

### Base editor profiles and activity windows

The base editor profile file describes the properties of each base editor. It specifies two editing windows to distinguish high-activity editing from detectable editing (Table 1).

**Table 1:**
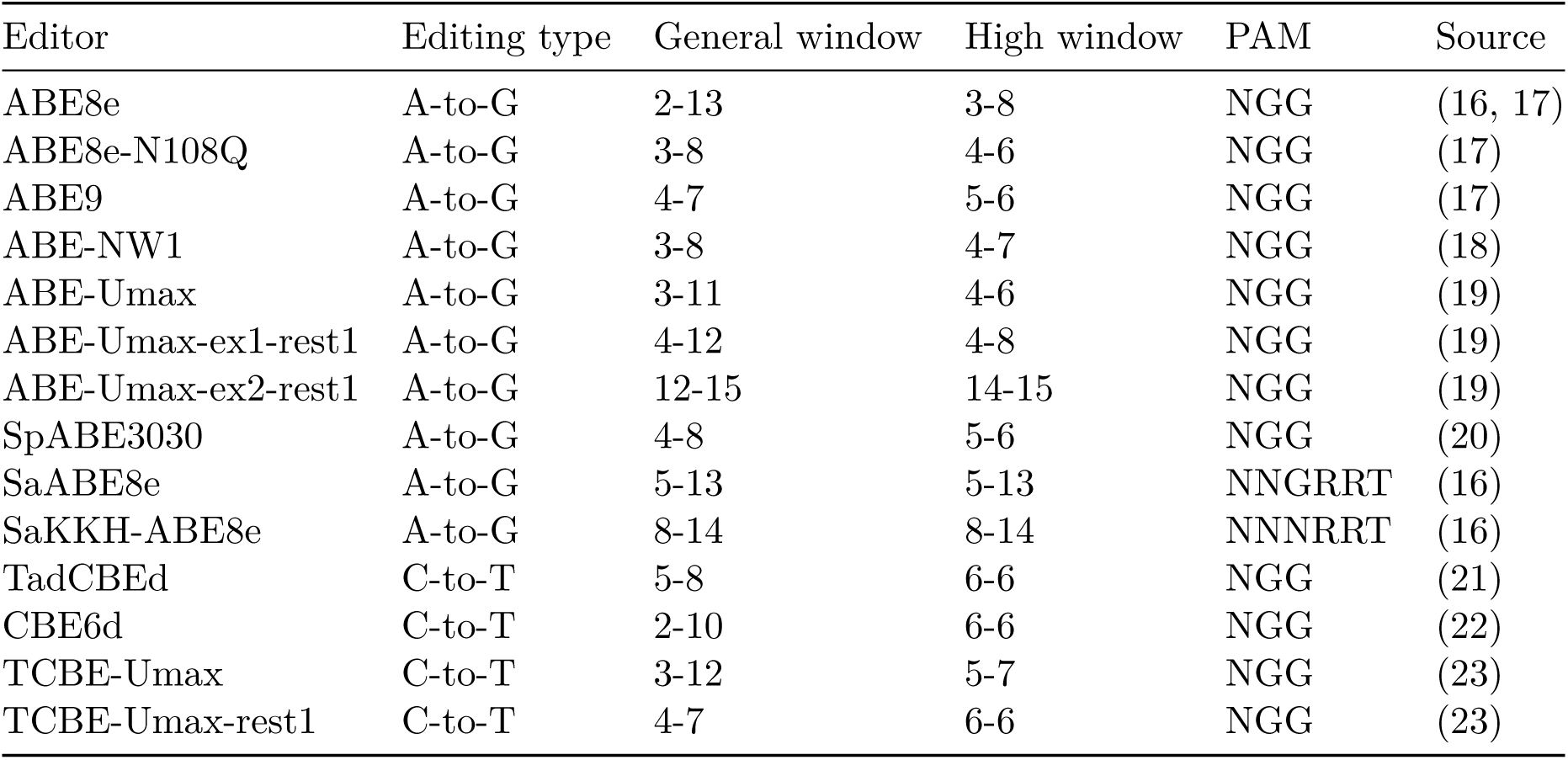
Simplified table of the base-editor profiles included in BEguided and BEguidedOFF.

#### High-activity window

The high_edit_start/high_edit_end coordinates define the guide sequence positions at which the editor converts the target base at ≥ ∼50% efficiency. These positions are used in the **Guidefinder** stage: a target is only reported as editable if the target base falls within this high-activity window.

#### General editing window

The general_edit_start/general_edit_end coordinates encompass positions in the guide sequence where editing is still detectable at ≥ ∼10% efficiency. This broader window is used for **bystander analysis** to identify all possible bystander edits and by **BEguidedOFF** to determine whether an off-target site can be edited by the defined base editors.

Preset parameters were curated from the primary publications of each editor, and only editors that achieved ≥ ∼50% editing efficiency in their source publication are included; editors whose general activity window is too broad for bystander-free design were likewise excluded. The source column records the exact reference for every profile.

For every canonical NGG SpCas9 profile, BEguided and BEguidedOFF additionally provide NGN-PAM (SpCas9-NG/SpG) and NNN-PAM (SpRY) versions — identical copies of the corresponding entries with only the PAM changed. Their activity windows are expected to be similar to those of the corresponding canonical profiles, although lower editing efficiencies should be anticipated because relaxed PAM variants generally exhibit reduced on-target activity compared with canonical NGG-targeting SpCas9 (24, 25). This derivation rests on the close similarity of the engineered Cas9 backbones to SpCas9 and is experimentally supported for ABE8e, which edits efficiently with SpCas9 and SpCas9-NG (16) and with SpRY (26); most other ABE profiles are themselves derived from, or closely related to, ABE8e, so their deaminase module is expected to behave similarly across backbones. For many of the remaining NGN- and NNN-PAM profiles, however, no editor-specific experimental evidence exists, and their tabulated activity windows are inherited assumptions rather than measured values.

### Input and output

#### Input files

The target file is a TSV or XLSX table of genomic coordinates (1-based), reference and alternate alleles, plus optional metadata (gene symbol, mutation ID). Additional required inputs are a reference genome (FASTA), gene annotation (GFF), and the user-extensible base-editor profile table described above. Optionally, a pre-computed BEguidedOFF genome encoding can be generated — a 2-bit .npz for the GPU-accelerated search or an FM-index (.fmi) for the CPU-based fast search.

#### Output

The final output contains one row per candidate guide and columns for guide sequence, strand, genomic coordinates, base editor profile, bystander positions, bystander mutation summary (protein-level consequences). When off-target search is performed, the columns off-target count, worst off-target CCTop score, worst off-target genomic location, and sequence of the worst off-target are also included.

### Guide RNA scoring

BEguided employs a lightweight on-target quality score called CRISPRater (15), which is used in the CCTop Cas9 target prediction tool (27). The score is computed from the 20-nt protospacer sequence alone and does not take the PAM into account. This keeps the score comparable across editor profiles with different PAM specificities, including the relaxed-PAM SpCas9 variants, but it also means that PAM-dependent differences in editing activity are not captured.

In addition to the guide score, two sequence-level properties are evaluated for every guide: the protospacer GC content and the presence of homopolymers. A GC content between 40% and 60% (inclusive) is classified as *high* and between 30% and 70% (inclusive) as *medium*. A run of four or more identical bases (homopolymer) is treated as unfavorable and rules the guide out of both the high and the medium class. Together with the guide score, these criteria form the joint on-target quality classification used throughout this study (see below).

Beyond these criteria, every ABE guide is additionally annotated with a *trinucleotide edit-context score*: the normalized ABE8e editing proportion for the trinucleotide flanking the target adenine (X-A-Y in mRNA sense, where X and Y denote the bases immediately upstream and downstream) (28). The score ranges from 0 to 1 (CAC = 1.0, AAA ≈ 0.02) and reflects how permissive the sequence context immediately surrounding the edit is. It can be applied as an optional filter (trinuc_edit_score_min); because the underlying table was derived from ABE8e data, it serves as an approximation for other ABE profiles, while CBE guides carry no score.

### BEguidedOFF: off-target prediction for base editors

BEguidedOFF was developed as a companion tool to BEguided and performs genome-wide off-target discovery for base editors. Keeping the search backend separate means its requirements (a compiled scanning kernel, a CUDA-capable GPU for the exhaustive search mode, and a one-time converted encoding of the reference genome) do not burden the core pipeline, and the search can be used, extended, or replaced independently of the guide-design logic.

Starting from a set of 20-nt guide RNAs and their base-editor profiles, BEguidedOFF scans both strands of the genome for all sites that carry the editor’s PAM *family*, i.e. the IUPAC motif defined in the profile (e.g. NGG for SpCas9, NNGRRT/NNNRRT for SaCas9, NGN for SpCas9-NG/SpG, NNN for SpRY), rather than the concrete on-target PAM, and lie within a user-defined mismatch tolerance of the guide, excluding the on-target locus itself. A distinctive feature is **editable-base-aware off-target filtering**: a site is reported only if it is *editable* (an A for adenine base editors or a C for cytosine base editors within the editor’s general editing window), not merely alignable. Each editable off-target is scored with the CCTop metric (see below), and the number of editable off-targets below a user-set score threshold is counted per guide. Guides are grouped by unique (sequence, PAM) pair so repeated sequences are scanned once.

BEguided calls the library with n_top_offtargets=1, so only the single worst off-target per guide is returned; these are the two values BEguided merges into its results table as worst_offtarget_score and offtarget_count (step 4, via the Python API). Run standalone on any guide set, the top-N parameter is user-settable (default 3) and all hits can be returned as a long-format table listing hit coordinates, sequence and PAM, mismatch positions, and scores. Two search modes, an exhaustive GPU mode and a fast CPU mode, are described below.

BEguidedOFF uses the CCTop metric (27) as its default off-target score. CCTop penalizes mismatches based on their distance from the PAM: proximal mismatches are weighted more heavily than distal ones via the formula Σ 1.2^p, where *p* is the mismatch position (1-indexed from the 5′ end of the 20 bp guide). The score is fast to compute, largely nuclease-agnostic, and interpretable: 0.0 indicates a perfect match, and higher scores correspond to progressively less concerning alignments. Guides with no detectable off-targets are left blank (NaN) and pass every score-based filter.

In its exhaustive search mode (extensive), the reference genome (GRCh38, Ensembl release 116) is encoded once into a compact 2-bit .npz format (∼1.3 GB including the pre-computed reverse complement) and scanned on a CUDA-capable GPU by brute force using XOR+popcount mismatch detection on 40-bit packed guide integers, with IUPAC PAM families encoded as 4-bit masks. The genome is processed in 200 Mbp chunks to bound GPU memory; guides that overflow the per-chunk hit buffer are automatically re-scanned with an escalating buffer so the returned hit list is exact, and guides that still overflow at the ceiling fail with an explicit error.

A second search mode (fast) performs the same discovery on a CPU via an FM-index (a compressed suffix array over the reference genome (29), the structure underlying mainstream short-read aligners (30)) and powers the public web application on GPU-less infrastructure. Instead of a mismatch tolerance, it takes a CCTop score cutoff (by default the guide-removal threshold) and exploits the penalty formula: because 1.2^p grows toward the PAM-proximal end, any site scoring below the cutoff must match the guide exactly in the 12 positions adjacent to the PAM (for the default cutoff of 5.0). The fast mode therefore locates this exact PAM-proximal anchor on both genome strands in the FM-index (reverse-strand sites via reverse-complement anchors), extracts the full protospacer-plus-PAM window, verifies the IUPAC PAM family, and scores the complete alignment; candidate generation is sound rather than heuristic, so no sub-cutoff site can be missed. The FM-index (∼1.7 GB for GRCh38, built once from the reference FASTA) is memory-mapped at query time, and results are streamed per guide so only the worst hits and the below-threshold count are kept; a cap guards against low-complexity anchors, and cutoffs so permissive that no exact region remains (≥ 1.2^20 ≈ 38.3) are rejected in favor of the exhaustive mode.

Users set three thresholds: max_mismatches (maximum mismatches for an off-target to be reported), count_threshold_score (CCTop upper bound for counting an off-target), and remove_guides_score (minimum worst_offtarget_score to retain a guide); in the fast mode, remove_guides_score doubles as the search cutoff, so the screen enumerates precisely the off-targets that could trigger removal.

BEguidedOFF was validated against Cas-OFFinder v2.4.1 (31) using the exhaustive GPU mode on 394 patient-mutation guides across the three PAM families (≤ 5 mismatches, ∼1.2 million off-target calls). A reference N inside the guide window is treated as a mismatch (strand-aware, matching Cas-OFFinder semantics; implemented via an N-run sidecar next to the FM-index). With these semantics, concordance is 100.0% for every PAM family and overall (1,243,721 unique off-target calls, zero differing calls), and the one repetitive guide that previously exceeded the hit buffer is enumerated exactly via the automatic re-scan. The two search modes were verified to be interchangeable for the removal decision: on 100 real mutation guides spanning all PAM families, and on a stress set of 24 highly repetitive guides (912 guide×editor rows) that exhausts the GPU hit buffer, both modes returned identical verdicts, identical sub-cutoff off-target sets, identical below-threshold counts, and identical top-3 lists. Because the fast mode restricts the search to sites that can influence the removal decision, it is roughly 12–31-fold faster than the exhaustive GPU scan, which is the property that makes interactive off-target screening of many targets across all 38 editor profiles feasible in the web application.

### Web implementation

BEguided is also available as a web application at https://beguided.buchholzlab.org/, a FastAPI server that executes the same pipeline with the reference genome and gene annotation preloaded, covering guide design, bystander annotation, filtering, and the optional off-target screen. Three input modes cover the typical use cases. In *sequence* mode, one or more target sequences are pasted with the desired edit marked inline (e.g. [A>G], as in Figure 2); the pasted sequence serves as the reference, so guide scoring runs on the user’s sequence, while bystander annotation, which requires a genomic transcript annotation, is not performed; results carry no genomic coordinates. In *single* mode, a target is entered manually by chromosome, 1-based position, and the desired base substitution. In *spreadsheet* mode, a table of targets (chromosome, position, target base, desired base) is uploaded and processed as one job. In all modes, checkboxes select all, a subset, or a single base-editor profile, and custom base-editor profiles can be defined directly in the interface.

Results are presented as an interactive table rather than a static list of all possible guides. Dynamic filters — bystander consequence, guide score, GC content, homopolymer status, and, for adenine base editors, the edit-context score — re-render the table instantly, with defaults set to the pipeline’s recommended guide criteria (synonymous or milder bystanders, acceptable guide score and GC content, homopolymers excluded). A summary bar chart counts, for each base editor, the targets still represented under the active filters, making the impact of editor choice and filter stringency immediately visible. Targets for which no guide was designed, or none matching the active filters, remain visible as placeholder rows rather than disappearing from view. A condense toggle merges guides that differ only in the base editor, and results can be downloaded as TSV or XLSX in the currently displayed state.

The public instance also includes an off-target screen: a *Check severe off-targets* button scans exactly the guides currently visible after filtering, using the fast FM-index mode of BEguidedOFF, which enumerates all editable sites with a CCTop score at or below the effective search cutoff (≈ 5.2 for the default settings, i.e. an exact 12-bp PAM-proximal anchor; the counting threshold of 15.0 is clamped to this cutoff), and reports for each guide the worst off-target found and a pass/fail verdict against the pipeline’s default removal thresholds: a guide fails if its worst off-target scores below 5.0, or if more than 100 editable off-targets are found. Only the worst hit and the count are returned rather than the complete hit list, so a “pass” verdict means that no off-target that severe was detected, not that the guide has no off-targets. The full genome-wide search with all settings is available through the command-line pipeline.

### Guide qualification criteria

All downstream analyses apply a uniform qualification scheme to the complete pipeline output, identically for the cBioPortal and ClinVar target sets. For the bystander criterion, the pipeline’s per-transcript bystander consequences are collapsed into four ordered severity tiers: no bystanders, synonymous, non-coding (target base or bystanders outside an annotated CDS), and nonsynonymous (abbreviated *nonsyn* in the figures; any amino-acid change, including stop-gain). Every guide with a valid PAM is evaluated against three criteria, which also provide the tier categories of the attrition funnels presented below; targets for which no guide can be designed are counted as not targetable. *Bystander risk* is rated by the mildest of a guide’s predicted bystander consequences: no bystanders are preferred over synonymous, followed by non-coding and nonsynonymous. *On-target quality* combines the guide score, protospacer GC content, and homopolymer status into three classes — high (guide score ≥ 0.75, GC content between 40% and 60% (inclusive), no homopolymer of four or more identical bases), medium (guide score ≥ 0.5, GC content between 30% and 70% (inclusive), no homopolymer, and not already high), and low (all remaining guides) — extending the GC-content and homopolymer classes defined above. *Off-target safety* is classified as high (off-target count < 50 and worst off-target score > 10), low (off-target count > 100 or worst off-target score < 6), or medium (all remaining guides). A guide is **qualified** only if it satisfies all three criteria simultaneously: no or synonymous bystanders, high or medium on-target quality, and high or medium off-target safety.

Guides are ranked with a pass-first ordering in which a guide that satisfies a quality criterion always outranks a guide that fails it, with remaining ties broken by the mildest bystander consequence, then the best on-target quality, then the safest off-target profile, and finally the highest guide score. The top-ranked guide represents each target in all summary statistics, so a target with at least one qualifying guide is always represented by a qualifying guide, making qualified counts identical to counting targets with at least one qualifying guide.

Targetability and qualification were quantified per editor panel: the percentage of targets that are targetable and the percentage of targets represented by a qualified guide. Attrition funnels (Sankey diagrams) trace each target through the four cascading stages of PAM availability, bystander consequence, on-target quality, and off-target safety, up to the qualified verdict, based on the representative guide. The attribution of targetable and qualified targets to Cas9 families was visualized as Venn diagrams over unique chromosome–position–strand targets, one set per family. Statistical analyses and visualizations were performed in R v4.6.1 (32) with the tidyverse v2.0.0 (33), ggsankey v0.0.99999 (34), and ggVennDiagram v1.5.7 (35) packages.

### cBioPortal cancer mutation analysis

cBioPortal is a public web resource for exploring, visualizing, and analyzing multidimensional cancer genomics and clinical data, integrating harmonized data from large cancer studies such as TCGA, MSK-IMPACT, MSK-CHORD, and CPTAC (36–38).

To quantify the global targetability of recurrent cancer mutations, we aggregated all somatic mutations — single-nucleotide variants (missense, nonsense, splice-site, and non-coding) as well as small insertions, deletions, and multi-nucleotide variants — from 38 curated public cBioPortal studies (TCGA pan-cancer cohorts, MSK-IMPACT, MSK-CHORD, and CPTAC; GENIE and other token-gated studies were excluded), accessed on 2026-10-31, lifting all variants to GRCh38. Mutations were classified as ABE-editable (A→G or T→C), CBE-editable (C→T or G→A), or not base-editable (all remaining single-nucleotide substitutions, which require conversions neither editor can do, and all insertions, deletions, and multi-nucleotide variants, which cannot be corrected by adenine or cytosine base editors). For the deep-dive analysis, we selected the 1,000 most frequently occurring ABE-editable mutations and the 1,000 most frequent CBE-editable mutations (ranked by aggregate sample counts across the contributing studies) and processed them with BEguided.

Both top-1,000 lists were processed with BEguided using all 38 built-in base-editor profiles (ABE and CBE profiles in separate runs) with BEguidedOFF off-target search enabled in the exhaustive GPU mode. No pre-filtering was applied (guide-score threshold 0; GC-content, homopolymer, and edit-context filters disabled), so the qualification scheme above was applied to the complete output of every profile. The off-target search used a mismatch tolerance of 5 and a counting threshold of 15.0, with guide removal disabled so that every guide remains represented and classifiable. Pre-computed guide sets for this analysis are provided in the accompanying dataset repository (see Data Availability).

### ClinVar inherited-disease variant analysis

To extend the analysis beyond somatic cancer mutations, we compiled an analogous target set from ClinVar (39), a public archive of relationships between human sequence variants and phenotypes as asserted by clinical laboratories. From the ClinVar variant summary release 2026-09 (GRCh38), we selected the 1,000 most frequently reported pathogenic or likely pathogenic single-nucleotide variants addressable by adenine or cytosine base editors, ranked by the number of independent submitter reports. The set comprises 879 germline and 121 germline/somatic variants. All variants were processed with BEguided and BEguidedOFF using the full base-editor panel and the same filtering and off-target settings as the cBioPortal analysis. Guides were ranked and variants classified as qualified as described above; because BEguided designs guides on both strands of a variant, ClinVar targets are aggregated per variant rather than per strand. The pre-computed guide sets for this ClinVar target list are provided in the accompanying dataset repository (see Data Availability).

## Results

### Global base-editability of cBioPortal cancer mutations

From the 1,218,505 unique somatic mutations aggregated across the 38 curated cBioPortal studies (1,122,100 single-nucleotide variants and 96,405 insertions, deletions, and multi-nucleotide variants), we classified all variants by the required base-editor conversions. We found that **54.5%** of all unique somatic cancer mutations are theoretically addressable with either an ABE or a CBE. Considering SNVs alone, the addressable fraction would be 59.2% (Table 2).

**Table 2:** Base-editor addressable mutations across the 38 curated cBioPortal studies.

| Editor type | Unique mutations | % of total |
|---|---|---|
| ABE (A→G / T→C) | 537,627 | **44.1%** |
| CBE (C→T / G→A) | 127,044 | **10.4%** |
| Not compatible* | 553,834 | **45.5%** |

This global survey demonstrates that base editors have broad but incomplete reach across the oncogenic landscape. Among SNVs, the catalog is dominated by C>T (270,229) and G>A (267,398) transitions, whereas the opposite direction, T>C (64,311) and A>G (62,733), is roughly four times rarer; because correcting the former requires T→C or A→G edits — precisely the conversions ABEs perform — the ABE-editable fraction is the larger one.

### Beyond ABE8e: multi-editor screening expands qualified access to the top 1,000 ABE-editable cancer mutations

While global editability tells us what is theoretically possible, a more practical question is how many of the *clinically most relevant* mutations can be targeted with high-quality guides. We therefore ran the top 1,000 most frequent ABE-editable cBioPortal mutations through BEguided. Even before any quality criteria are considered, the choice of Cas9 backbone determines which of these targets can be reached at all (Figure 3).

**Figure 3:**
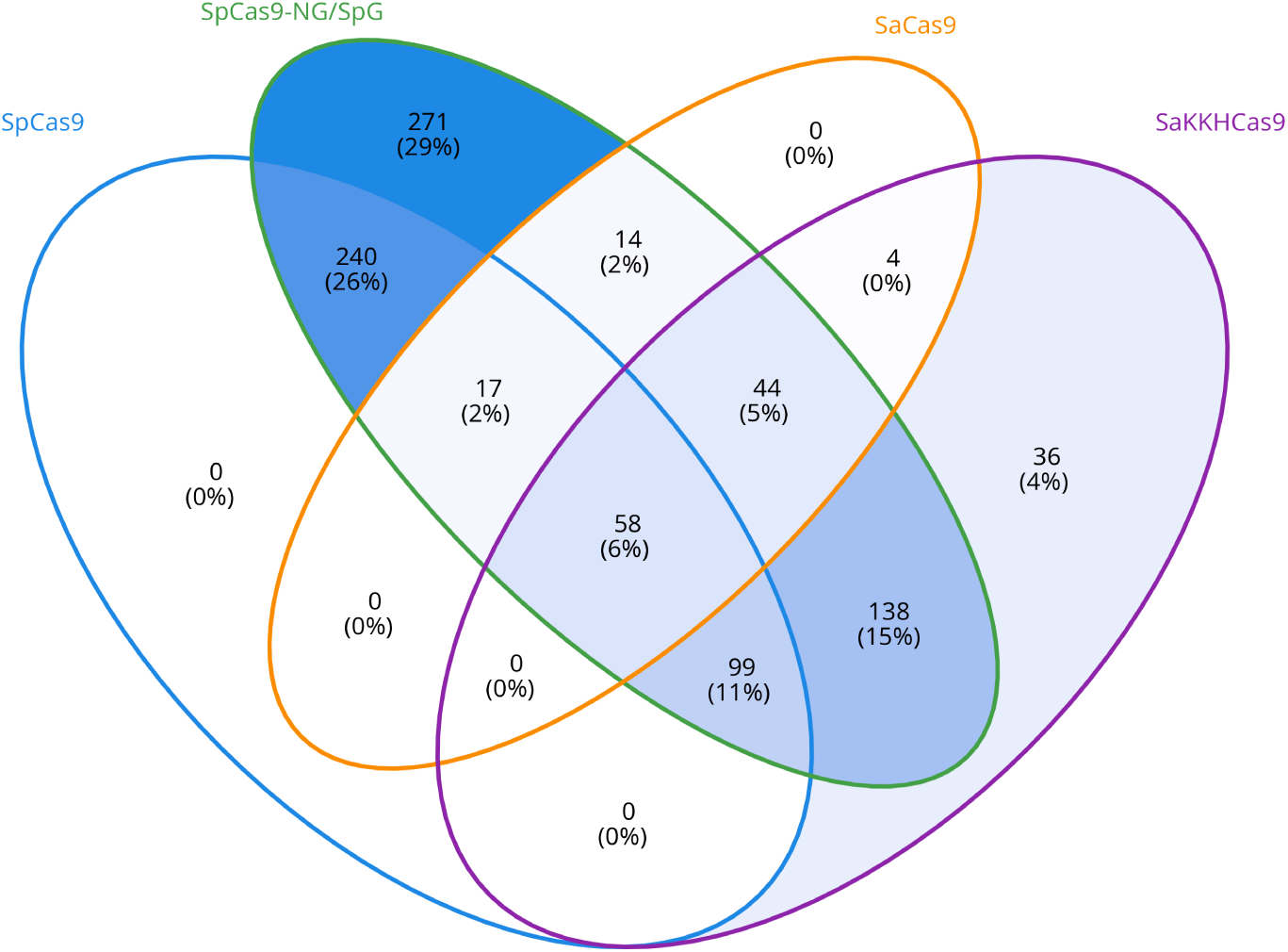
PAM-restricted targetability of the top-1,000 ABE-editable cBioPortal mutations across Cas9 families. Four-set Venn diagram of the 921 target mutations (unique chromosome–position–strand combinations) for which at least one guide RNA from any of the four non-SpRY Cas9 families places the target base within the editing distance of the corresponding adenine base editors. Sets: SpCas9 (NGG), SpCas9-NG/SpG (NGN), SaCas9 (NNGRRT), and SaKKHCas9 (NNNRRT). Numbers give target counts and percentages of the 921 targetable mutations.

Typically, when designing guide sequences manually, the focus is on one single well-characterized editor such as ABE8e (SpCas9). We first asked how many of these mutations yield a qualified guide when only ABE8e is considered. Of the 1,000 mutations, 329 (32.9%) are targetable, but only 7 (0.7%) of these have a qualified guide (Figure 8; Figure 4) — no or synonymous bystanders, high or medium on-target quality, and high or medium off-target safety (see Methods for details). This low yield underscores the restrictive nature of the SpCas9 NGG PAM combined with a relatively wide base editing window of TadA-8e, which results in bystander edits.

**Figure 4:**
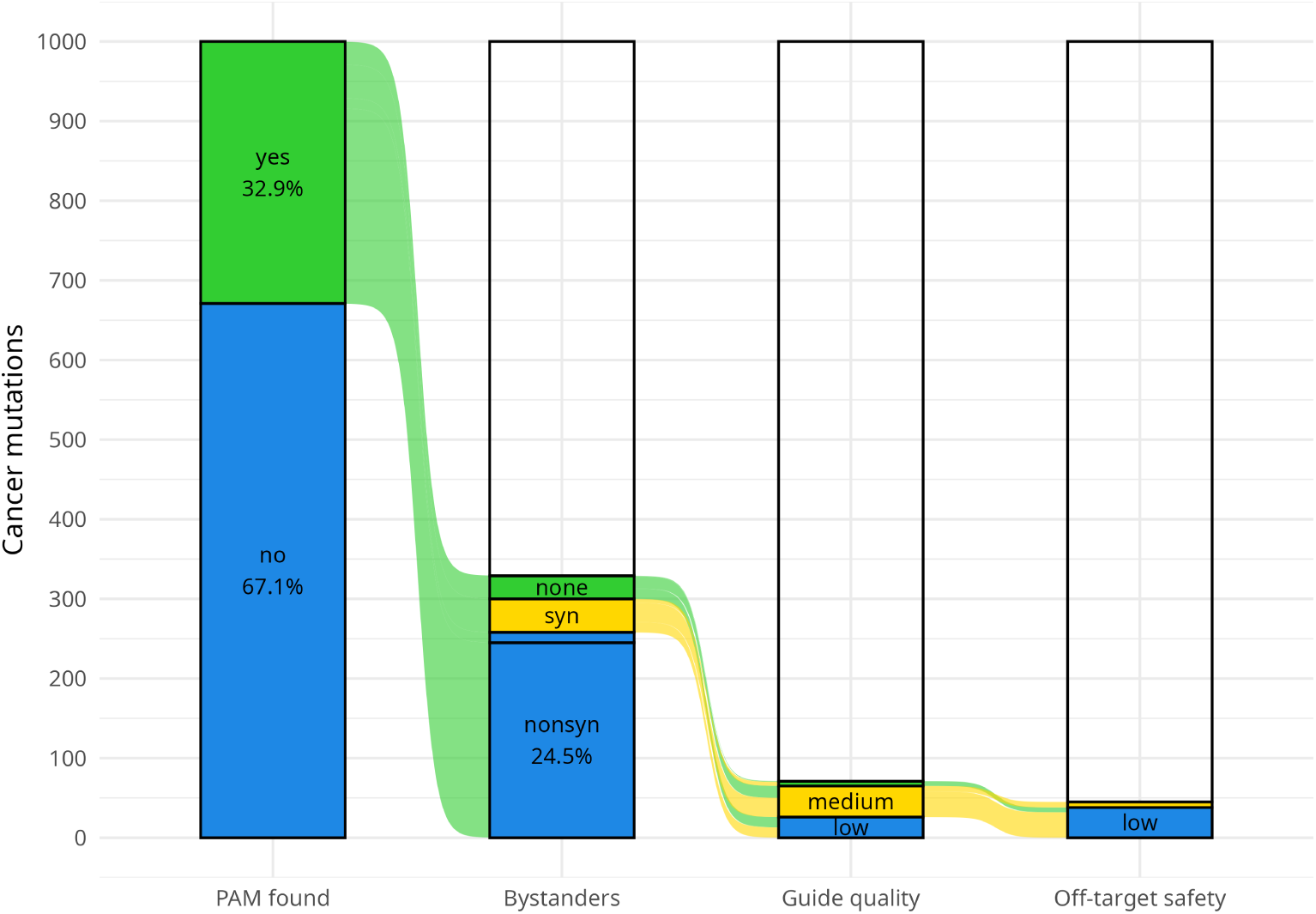
Attrition across filtering stages for the ABE8e single-editor panel. The total entering the funnel is 1,000 selected target mutations. Colors indicate favorable (green), intermediate (yellow), or unfavorable (blue) outcomes.

When the same targets are screened against the full panel of SpCas9-compatible adenine base editors in addition to ABE8e (ABE-NW1, ABE-Umax and derivatives, ABE9, and SpABE3030), the targetable fraction rises modestly to 414 (41.4%), but the fraction of qualified guides increases more than four-fold to 30 (3.0%). Strikingly, the entire targetability gain is contributed by a single profile, ABE-Umax-ex2-rest1: its editing window (guide positions 12–15) sits at the PAM-distal end of the protospacer, outside the PAM-proximal windows shared by all other SpCas9-compatible ABEs, making 77 target bases reachable that no other SpCas9-compatible ABE reaches. The gain in qualified guides, by contrast, is driven by the variety of editing windows, which give each target more editor options and thereby increase the fraction of targets that can be edited without bystanders (Figure 4; Supplementary Figure S1).

If one then also considers base editors with the Cas9 ortholog SaCas9, the number of targetable mutations increases to 650, and when considering guide quality, bystanders, and off-targets, the number of guides ends up at 100, more than doubling the SpCas9 ABE number (Figure 5; Supplementary Figure S2).

**Figure 5:**
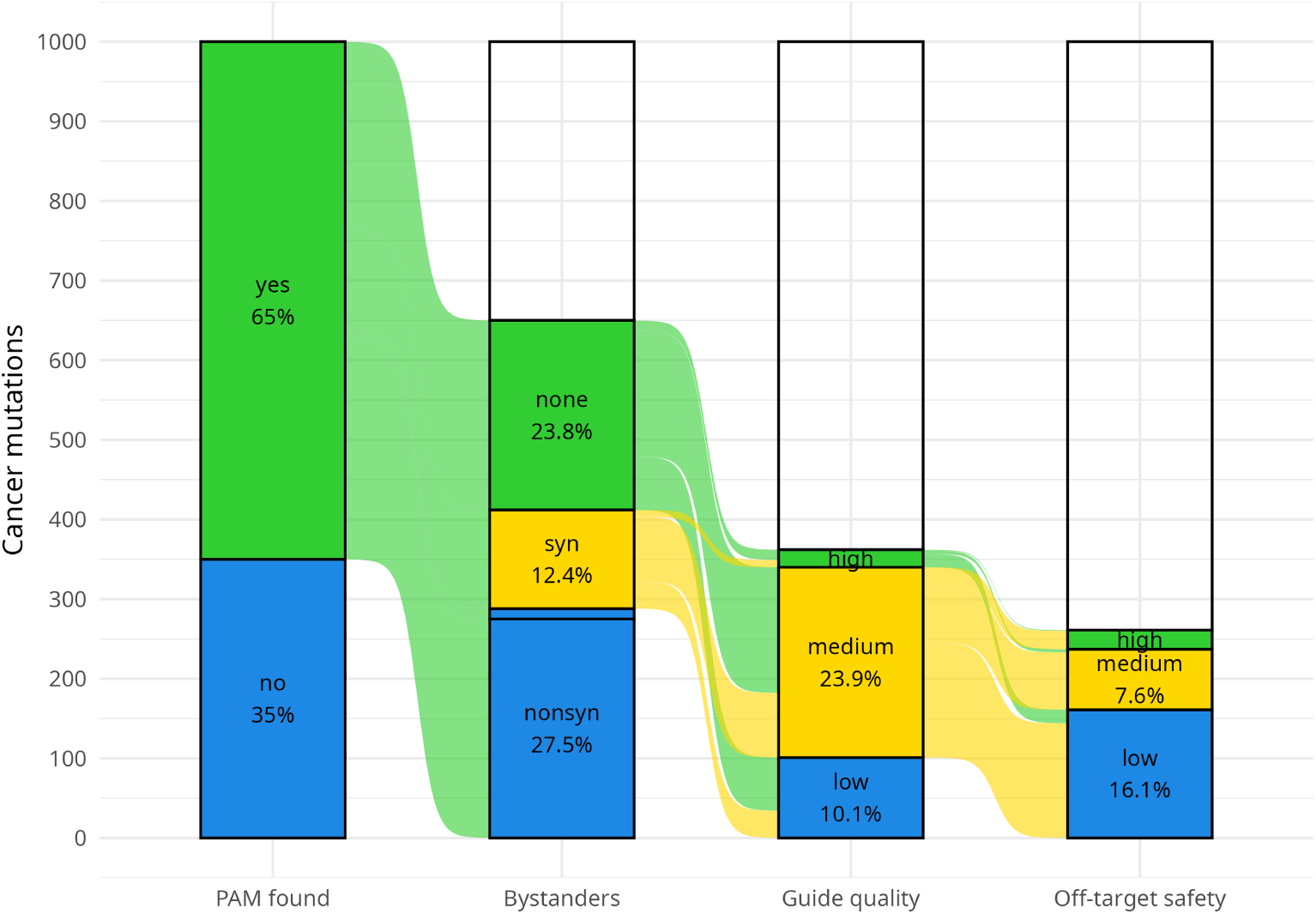
Attrition across filtering stages for the combined SaCas9 + SpCas9 ABE panel. The total entering the funnel is 1,000 selected target mutations. Colors indicate favorable (green), intermediate (yellow), or unfavorable (blue) outcomes. Analogous attrition panels for all remaining editor combinations are provided in the Supplementary Materials.

Further increases can be achieved with the relaxed-specificity NGN-PAM SpCas9 variants SpCas9-NG and SpG (24), raising the panel to 881 targetable mutations and 96 qualified guides. The price of the expanded PAM repertoire is a higher off-target load: the SpCas9-NG/SpG panel reports substantially more off-target hits per guide than the canonical SpCas9 or SaCas9 panels (Supplementary Figure S3; Supplementary Figure S4; Supplementary Figure S5), because relaxed PAM recognition motifs are markedly more permissive than NGG or NNGRRT.

We also assessed a panel consisting only of SpRY-based ABEs, which recognize the maximally relaxed NNN PAM. Strictly, SpRY is most active on NRN PAMs (R = A or G) and retains lower but substantial activity on NYN PAMs (Y = C or T); because these two motifs together exhaust NNN, we assign all SpRY profiles the full NNN PAM without distinguishing the two classes (24). This leads to targetability of all 1,000 top ABE-editable mutations (100%), i.e. every target that yields any valid guide in our analysis. SpRY also delivers a competitive proportion of qualified guides: 19.1% of targets met all quality criteria (Figure 8; Supplementary Figure S5), a figure substantially higher than the SaCas9 + SpCas9 combined panel (10.0%) and somewhat higher than the ensemble that excludes SpRY (15.6%). The attrition funnel for SpRY (Supplementary Figure S5) shows that, while no mutations are lost at the PAM stage, the subsequent filtering stages (bystander analysis, on-target quality, and especially off-target scoring) remove many candidates. The NNN PAM reports vastly more off-target hits per guide than any other panel, and many of these off-targets carry concerning CCTop scores. Despite this off-target burden, SpRY outperforms every non-SpRY panel in the proportion of qualified guides, as no other panel matches its PAM coverage.

Combining SpRY with the SaCas9 + SpCas9 + SpCas9-NG/SpG panel (i.e. including every base-editor profile evaluated here) yields a targetable fraction of 100% (1,000/1,000) and a qualified guide fraction of 23.8% (238/1,000), the highest qualified-guide yield of any panel we evaluated (Figure 8; Figure 6). Adding SpRY to the combined panel therefore provides a meaningful absolute and relative gain in the proportion of high-quality guides, despite the additional off-target burden it imposes. Figure 7 attributes the 238 qualified targets obtained with all ABE profiles to the Cas9 families and shows that a substantial fraction of them does not require maximally relaxed PAMs: 100 targets (42.0%) are covered by the canonical-PAM families alone — 30 by NGG-targeting SpCas9 editors and 74 by the SaCas9-backbone editors (SaCas9, NNGRRT PAM; SaKKHCas9, NNNRRT PAM). The sets are strictly nested — every target qualified by canonical SpCas9 is also qualified by SpCas9-NG/SpG, and both sets lie entirely within SpRY — so the relaxed-PAM families serve purely to extend reach, and canonical-PAM editors remain the preferable choice wherever they qualify a target (see Discussion). Conversely, the SaCas9 families qualify 79 targets, including 62 that no SpCas9-backbone family reaches and 47 that are not qualified by SpRY either; for the 32 targets qualified by both Sa-family and SpRY editors, the more restrictive SaCas9 PAMs are predicted to offer a safer alternative to SpRY, consistent with the higher off-target burden of relaxed PAM recognition (Supplementary Figure S3; Supplementary Figure S4; Supplementary Figure S5).

**Figure 6:**
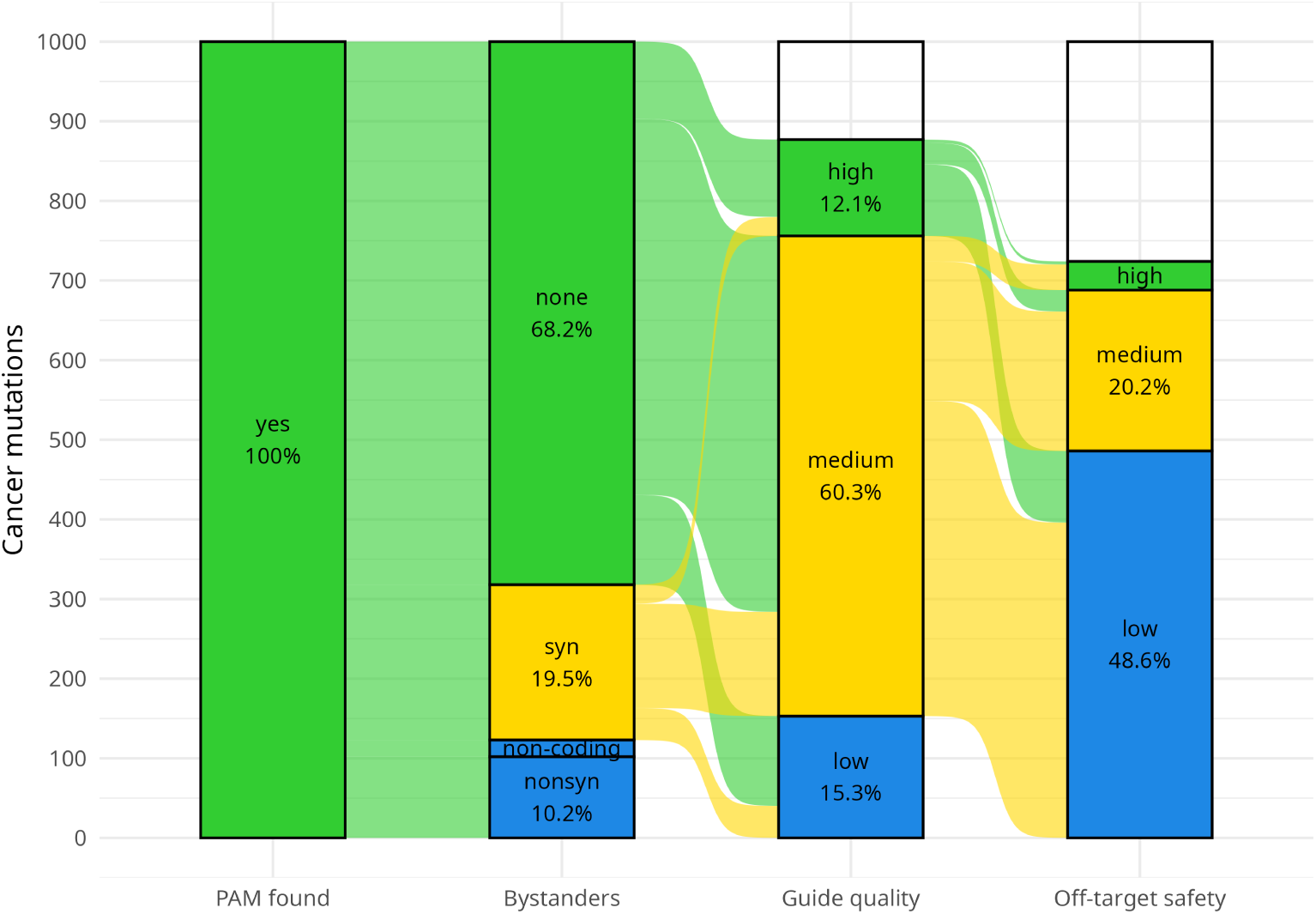
Attrition across filtering stages for the full ABE panel (SaCas9 + SpCas9 + SpCas9-NG/SpG + SpRY). The total entering the funnel is 1,000 selected target mutations, of which 23.8% yield a qualified guide. Colors indicate favorable (green), intermediate (yellow), or unfavorable (blue) outcomes.

**Figure 7:**
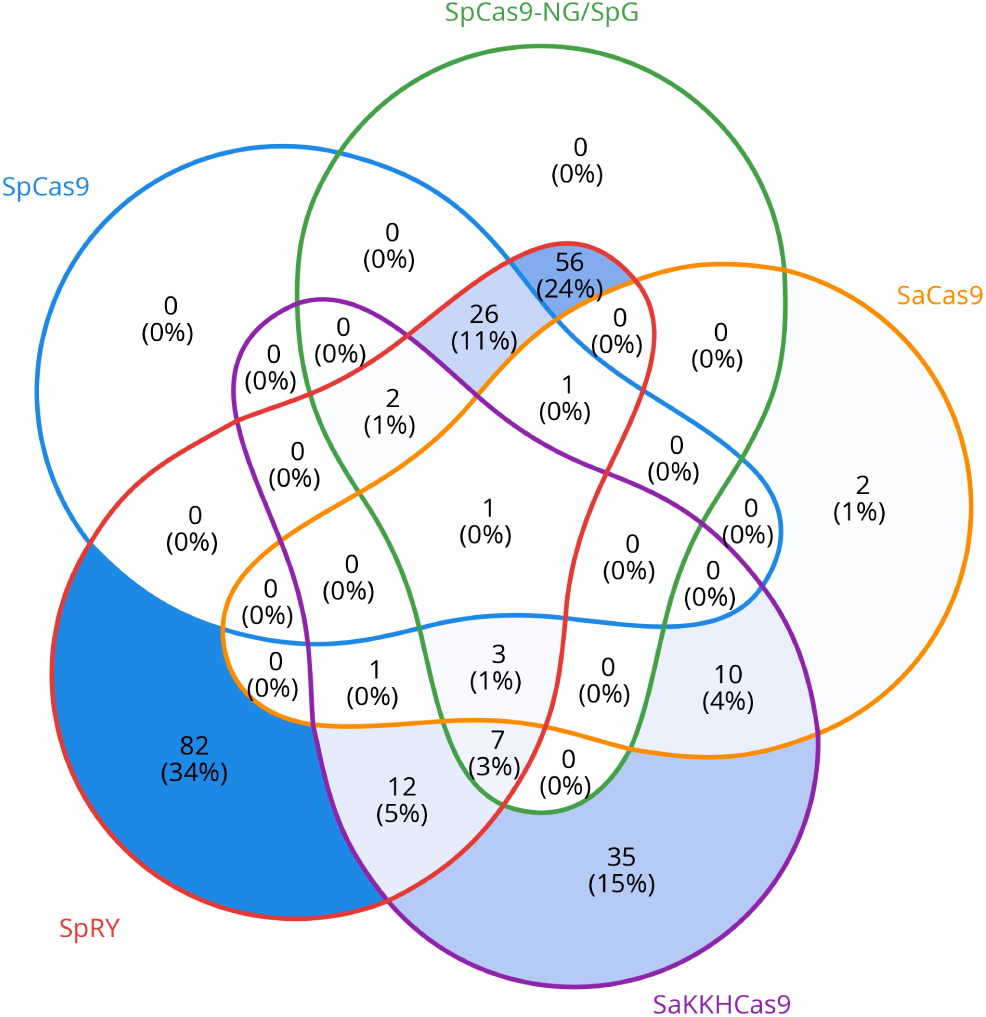
Targets with a qualified guide across Cas9 families, including SpRY. Five-set Venn diagram of the 238 ABE-editable cBioPortal targets for which at least one Cas9 family delivers a qualified guide (no or synonymous bystanders, high/medium on-target quality, and high/medium off-target safety). Each set counts the targets that yield a qualified guide when only that family’s base editors are considered: SpCas9, SpCas9-NG/SpG, SaCas9, SaKKHCas9, and SpRY (NNN PAM). Numbers give target counts and percentages of the 238 targets. The union of the five sets equals the qualified-guide count of the full ABE panel (Figure 8; Figure 6), since a target counts as qualified whenever any of its guides satisfies all criteria.

### A narrow window: cytosine base editors deliver few qualified guides for the top 1,000 CBE-editable cancer mutations

For completeness, we performed the same analysis for the 1,000 most frequent CBE-editable mutations, even though the edits CBEs perform (C→T and G→A) correct only the rare transition classes (T>C and A>G; together 127,044 SNVs, 10.4% of the catalog). CBE targetability is markedly lower than ABE targetability (Supplementary Figure S6): SpCas9 CBEs alone reach only 19.5% of the top targets, whereas SpCas9-NG/SpG CBEs expand coverage to 59.4%. The compressed CBE panel contributes to this picture: only four canonical SpCas9 CBE profiles (TadCBEd, CBE6d, TCBE-Umax, and TCBE-Umax-rest1) passed the inclusion criteria, because many published cytosine editors fall short of the activity threshold or carry editing windows too wide for bystander-free design.

More importantly, the four profiles are largely redundant: three share the same single-position high-activity window (position 6), reach an identical set of 85 targets, and add nothing to one another, while TCBE-Umax (high-activity window 5–7) subsumes all of them and single-handedly accounts for the panel’s full reach of 190 targets. Where the ABE panel gained reach through the genuinely new, PAM-distal window of ABE-Umax-ex2-rest1, the CBE panel’s reach is set by one profile’s slightly wider window.

The proportion of qualified CBE guides remains correspondingly modest (0.3–1.5% for the PAM-restricted panels, 6.6% when SpRY is included), and the window structure explains much of this attrition (Supplementary Figure S7; Supplementary Figure S8). Because bystander analysis is performed over the general editing window, only 4.7% (CBE6d, general window 2–10) and 7.8% (TCBE-Umax, general window 3–12) of their guides are bystander-free, compared with 38.8% (TadCBEd, general window 5–8) and 36.5% (TCBE-Umax-rest1, general window 4–7). The profile that extends targetability is thus precisely the one whose guides are most often compromised by bystanders, and — unlike the ABE panel, whose narrow distal-window profile ABE-Umaxex2-rest1 combines reach with bystander-free design — no CBE profile combines both properties. The remaining losses occur at the on-target quality and off-target stages, as in the ABE panels (Supplementary Figure S9; Supplementary Figure S10; Supplementary Figure S11).

### Beyond cancer: inherited pathogenic variants in ClinVar

The same workflow applies unchanged to germline variants that are candidates for correction rather than somatic lesions to be characterized. We therefore processed the 1,000 most frequently reported pathogenic and likely pathogenic ClinVar SNVs (39), comprising 879 germline and 121 germline/somatic variants, through the full pipeline, evaluating each Cas9 backbone with all of its adenine and cytosine base-editor profiles combined (Figure 9). ABE8e alone yields a qualified guide for only 6 of the 1,000 SNVs. Considering all SpCas9-compatible base editors (ABE and CBE) raises this to 35, adding SaCas9 to 89, the relaxed-PAM SpCas9-NG/SpG variants to 135, and SpRY to 206 — a roughly 34-fold increase relative to ABE8e that underlines the practical value of multi-editor screening for therapeutic variant correction. Of the 206 SNVs with a qualified guide, 194 come from the ABE profiles, which on their own target 818 of the 1,000; the CBE profiles cover the remaining 182 and contribute the other 12, so the two disjoint classes together account for all 1,000 targetable SNVs. This ABE-heavy split is consistent with the strong enrichment of pathogenic SNVs for transitions. The full panel’s 100% targetability is by construction: the SNV set was preselected for base-editor addressability, and the SpRY profiles recognize any PAM (NNN), so no site lacks a PAM.

**Figure 8:**
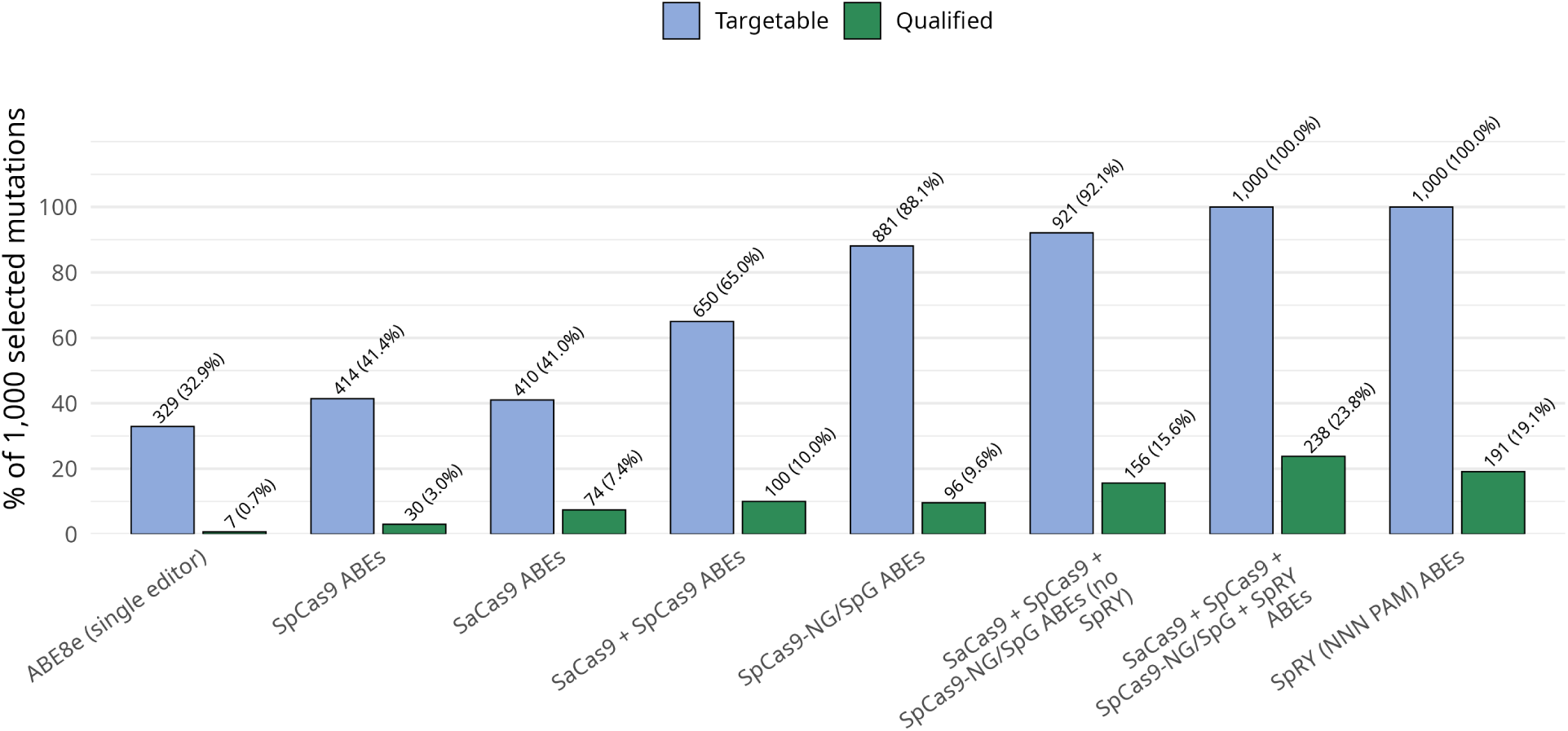
Targetability of the top-1,000 most frequent ABE-editable cBioPortal mutations across editor panels. Bars show the percentage of the 1,000 selected mutations that are targetable (at least one guide places the target base within the editor’s high-activity window) and the percentage of targets with at least one qualified guide — no or synonymous bystanders, high or medium on-target quality, and high or medium off-target safety (see Methods). Labels above the bars give target counts and percentages.

**Figure 9:**
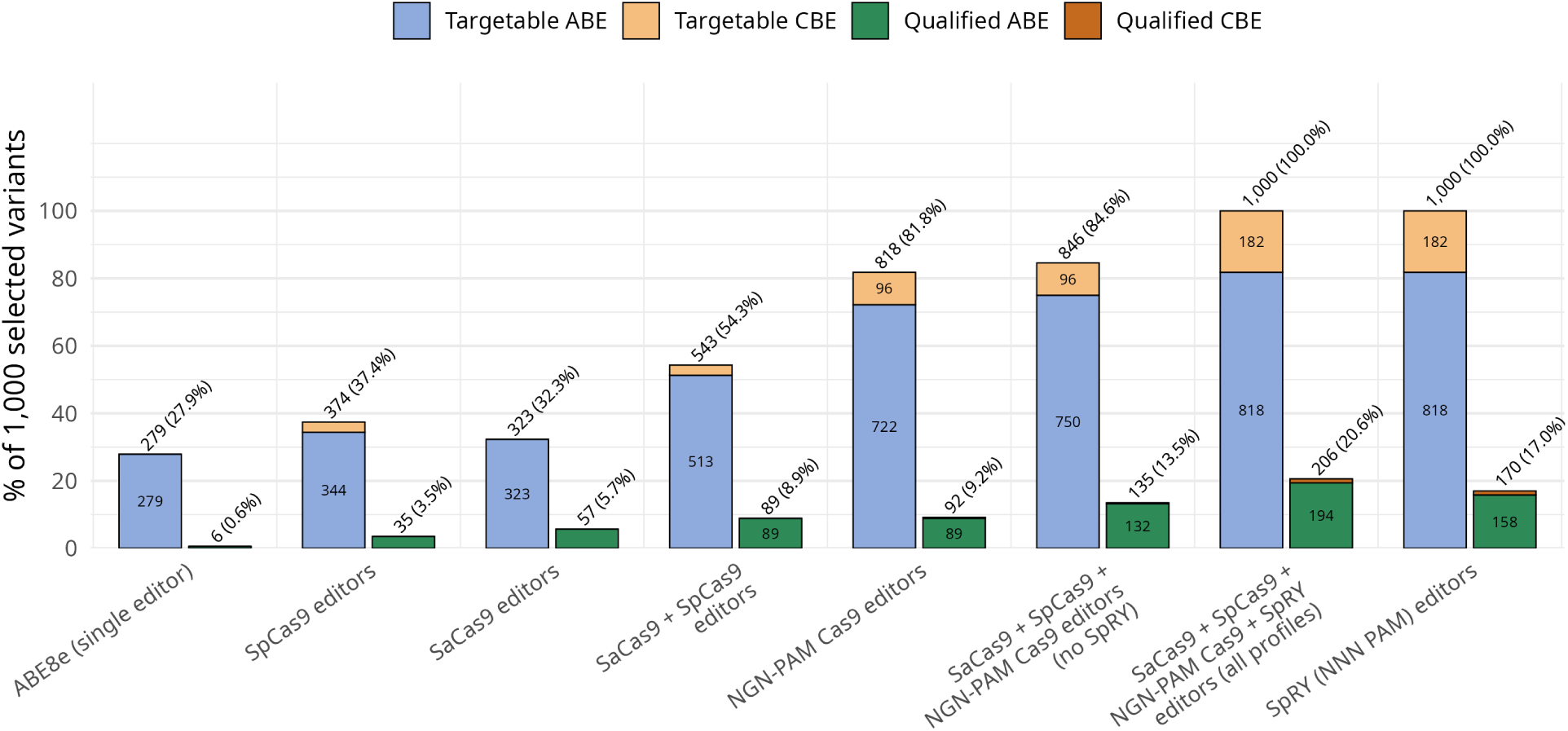
Targetability of the 1,000 most frequently reported pathogenic ClinVar variants across editor panels. Each panel combines all built-in base-editor profiles of the listed Cas9 backbone(s) — adenine- and cytosine-deaminase fusions together; the SaCas9 + SpCas9 + NGN-PAM Cas9 + SpRY panel therefore corresponds to all 38 editor profiles. Bars show the percentage of the 1,000 selected variants that are targetable and the percentage of variants with at least one qualified guide, as defined for Figure 8. Each bar is stacked by editor class: segments give the variants addressable by adenine base editors (blue/green) versus cytosine base editors (orange); the classes are disjoint, because A→G/T→C and C→T/G→A corrections require different editor types. Labels above the bars give variant counts and percentages.

## Discussion

### Design philosophy

BEguided was written for a specific mode of work: the user requires one or more defined single-base changes and needs a quick, broad check of the base-editing options available for those targets. For each edit, the pipeline asks which editor profiles can place a guide with a valid PAM over the target base and whether the resulting guides meet its quality criteria, rather than enumerating every guide that a given sequence could yield. Keeping the question this narrow keeps the output small, interpretable, and directly comparable across editor panels.

Web tools such as BE-Designer (12) and ACEofBASEs (13) take the opposite, sequence-centered paradigm: a reference genome is selected for off-target search, a DNA sequence or gene identifier is provided as input, and all potential base-editing guides for that sequence are enumerated. Such outputs tend to be large and hard to interpret, with few options to filter or rank candidates. BEguided instead inverts the input model: the user specifies the base and position to be edited and the base-editor profiles of interest (all, a subset, or a custom profile) and receives a pre-evaluated, filterable table focused on that edit, which streamlines guide design for targeted base editing. Because all editor–guide combinations for a target are evaluated in a single pass, the best available combination can be identified directly from the output — at the high throughput demonstrated by the 1,000-mutation cancer and ClinVar panels analyzed in this study.

### Editing without custom reference genomes

BEguided constructs guide sequences by inserting the desired target base directly into the extracted genomic context, even when the reference genome carries the wild-type allele at that position. The web tools discussed above cannot do the same: BE-Designer and ACEofBASEs enumerate guides for the sequence they are given, so corrective design against a mutant allele requires a modified reference genome for every individual SNV. BEscreen (14), which does accept specific variant coordinates, has the opposite limitation: it only supports variants that convert the existing reference base into another base, so the corrective direction — converting a mutant, non-reference base back into the reference base — cannot be specified. BEguided therefore designs corrective guides straight from the unmodified reference, without custom reference genomes.

The trade-off is that nearby polymorphisms in the target genome are not taken into account when extracting protospacer context. If the clinical sample or cell line carries additional SNPs within the 20-bp guide or PAM region, binding properties may deviate from the in silico prediction. Consequently, BEguided is best understood as a rapid first-pass screening instrument that identifies which mutations are theoretically targetable and which editor configurations offer the cleanest profiles. For individual therapeutic designs, researchers should verify guide sequences against the exact patient or cell-line genome and supplement BEguided output with experimentally validated on-target efficiency predictors.

### Complementary tools

BEguided and BEguidedOFF are intended for screening and triage, not for replacing specialized predictors. For applications that demand the highest predictive fidelity, several complementary tools can be applied to the short-list of candidate guides:

- **On-target efficiency.** Machine-learning models such as BE-Hive (40), BEDICT2.0 (25), BEdeepon (41), DeepBE (42), and CRISPRon/off (43) provide experimentally trained predictions of base-editing efficiency. For canonical SpCas9 NGG contexts, Rule Set 3 (44) offers an additional layer of activity prediction. These tools can be applied to BEguided’s candidate short-list to rank predicted on-target performance.
- **Off-target discovery with indels.** BEguidedOFF performs mismatch-only alignment. Cas-OFFinder with bulge settings (or similar tools) can supplement these results for contexts where insertion or deletion off-targets are a concern. BEguided’s plain TSV format makes this hand-off straightforward.

### Limits of the current evidence for editor recommendation

Throughout this study, BEguided has been presented as a comparison framework rather than a recommendation engine. The pipeline reports how each editor profile performs on a given target set; it does not rank editors or predict which variant will edit most efficiently in a particular experiment. This restraint reflects the current state of the field. Many base-editor variants have been characterized only in the publication that introduced them and on a small number of targets, so their reported efficiencies and the editing windows derived from those reports rest on limited evidence. Editing efficiency is further modulated by cellular context, including cell type, chromatin accessibility, and the transcriptional state of the locus (40, 42); a window calibrated in one system need not transfer to another.

Comprehensive, cross-editor benchmarking efforts such as BE-Hive (40) and BEDICT2.0 (25) are important steps toward filling this gap, but their editor panels do not yet cover every variant or the most recent enzymes. In the absence of such data, an explicit recommendation between, for example, ABE-Umax (19) and ABE9 (17) cannot be made on evidence. The defensible statements are narrower. The extensively validated ABE8e (16) remains the safest default — a judgment based on the breadth of independent validation rather than on head-to-head efficiency comparisons — and relaxed-PAM variants such as SpCas9-NG, SpG, and SpRY (24, 45) generally trade reduced on-target activity for an expanded targeting range and a higher off-target burden, although the magnitude varies by locus and study. The BEDICT2.0 benchmark quantifies this trade-off for base editing itself: average A-to-G editing with ABE8e on each backbone’s preferred PAM class fell from 64.9% (SpCas9, NGG) to 34.2% (SpG, NGN) and 33.9% (SpRY, NRN), and SpRY-ABE8e reached NYN motifs only to a lesser extent (25). The NNN-PAM profiles used in this study therefore describe targeting reach rather than uniform expected editing efficiency.

For these reasons, base-editor profiles, including their editing windows, were designed as an open, user-editable input. A laboratory can encode its own measurements and immediately obtain recommendations tailored to its experimental system without modifying the pipeline. Any resulting ordering of editors then reflects that laboratory’s own data rather than assumptions built into the tool, which keeps the tool adaptable as new data and newer variants emerge.

### Beyond SNV correction

The survey in this study was primarily framed around the correction of nucleotide variants in a disease context such as cancer, but the constraints BEguided weighs (PAM availability, editing-window placement, bystander consequences, guide quality, and off-target burden) are identical for any single-nucleotide engineering goal, and the pipeline accepts arbitrary reference genomes (FASTA + GFF) and target lists. Two applications follow directly. The first is the deliberate introduction of SNVs into a wild-type background — modeling a disease-associated allele or any other variant of interest in a model organism, cell line, or organoid. Bystander annotation remains essential: the intended edit must not be accompanied by an unintended coding change, and candidate guides must ultimately be re-validated against the actual model haplotype.

The second is the humanization of mouse cells. Genetically engineered mouse models are most informative when they carry the human-relevant allele, and H2M (46) predicts mouse variants that mirror the sequence and functional effects of human variants, providing base-editing libraries for 4,612 such mutations. Because BEguided runs on any annotated reference genome, the mouse-side variants from H2M can be processed directly against the mouse genome to compare all 38 editor profiles, evaluate bystanders on the mouse transcript annotation, and quantify off-target burden. This multi-editor evaluation goes beyond the single-PAM designs provided there and extends to humanizing loci outside the pre-computed H2M database.

Across our surveys, editability emerged not as a fixed property of a genomic position but as a function of the editor panel applied to it — the premise BEguided was built around. The pipeline turns a list of target edits into a side-by-side, bystander- and off-target-aware comparison of every built-in editor profile, extensible to user-defined editors, while keeping its predictions interpretable. We expect routine multi-editor screening to become the default first step of base-editing design, and we provide BEguided, BEguidedOFF, and the accompanying pre-computed guide sets so that taking it is a tool call rather than a manual design effort or a bespoke scripting project.

## Data Availability

The mutation data analyzed in this study are publicly available from cBioPortal (https://www.cbioportal.org/) and ClinVar (https://www.ncbi.nlm.nih.gov/clinvar/). The code and data underlying this work are available on GitHub:

- *BEguided* pipeline: https://github.com/ltschmitt/BEguided
- *BEguidedOFF* off-target library: https://github.com/ltschmitt/BEguidedOFF
- Pre-computed guide sets, processing scripts, and figure-generation scripts for the ClinVar and cBioPortal analyses: https://github.com/ltschmitt/BEguided-datasets

## Acknowledgements

During the preparation of this work, the authors used a range of open-weight large language models, applied through the OpenCode coding agent, to assist in drafting and editing the manuscript, developing the pipeline and analysis code, and generating the vector graphics of the figures as SVG code — the figures were authored as code, not produced by image-generation models. All AI-assisted content was reviewed and verified by the authors, who take full responsibility for the work. We thank ScaDS.AI Dresden/Leipzig and the Zentrum für Informationsdienste und Hochleistungsrechnen (ZIH) of TU Dresden for providing access to these models, and ZIH for hosting the BEguided web service.

## Author contributions

**Lukas Theo Schmitt:** Conceptualization, Methodology, Software, Investigation, Data Curation, Formal Analysis, Visualization, Project Administration, Writing – Original Draft, Writing – Review & Editing. **Pascal Wang:** Conceptualization, Software, Validation, Writing – Review & Editing. **Sara Orugo Medina:** Software, Validation, Writing – Review & Editing. **Moustafa Abohawya:** Software, Validation, Writing – Review & Editing. **Frank Buchholz:** Conceptualization, Funding Acquisition, Supervision, Writing – Review & Editing. **Shady Sayed:** Conceptualization, Data Curation, Funding Acquisition, Supervision, Writing – Review & Editing.

## Supplementary material

**Supplementary Figure S1:**
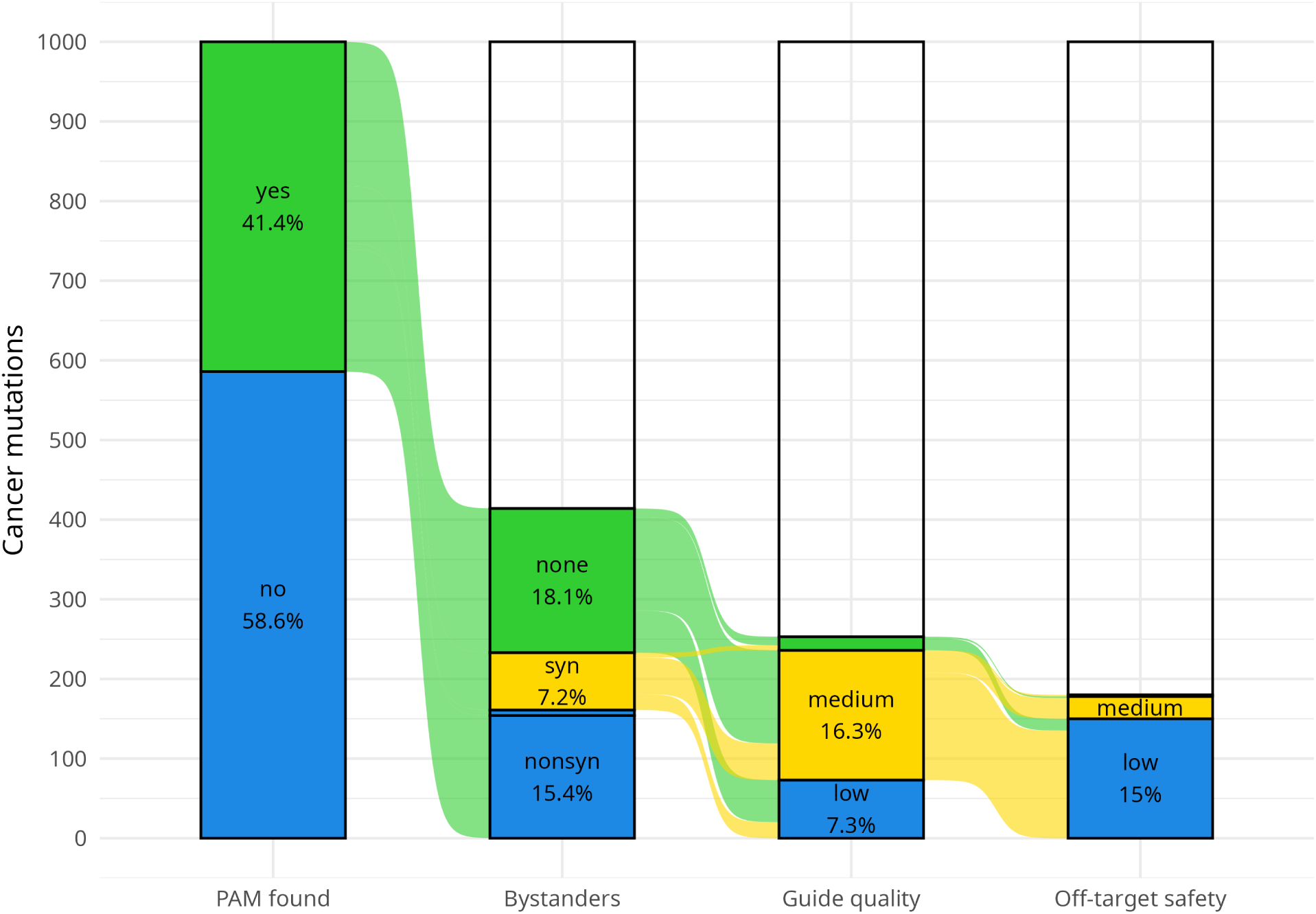
SpCas9 ABE attrition funnel for the top-1,000 ABE-editable cBioPortal mutations.

**Supplementary Figure S2:**
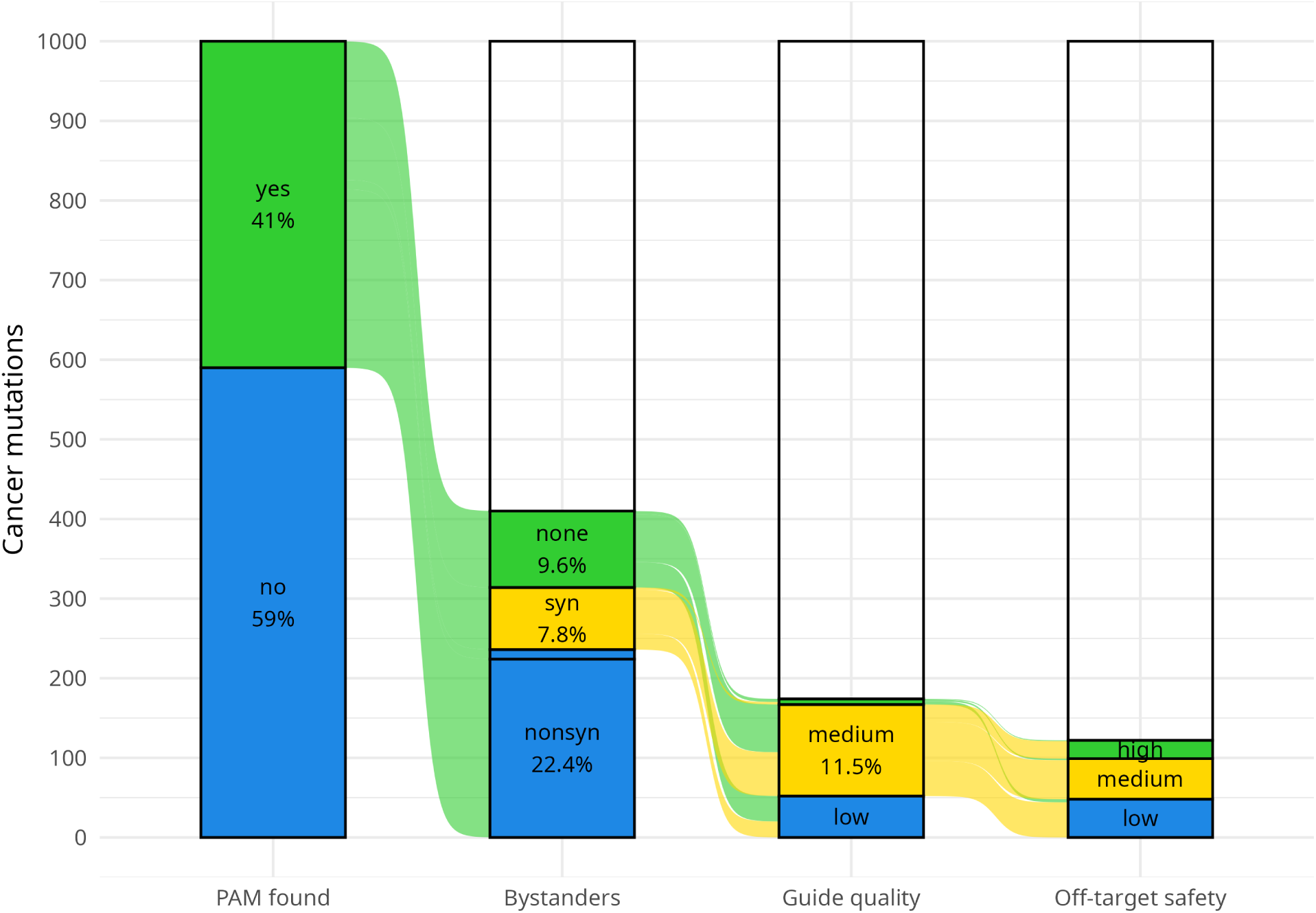
SaCas9 ABE attrition funnel for the top-1,000 ABE-editable cBioPortal mutations.

**Supplementary Figure S3:**
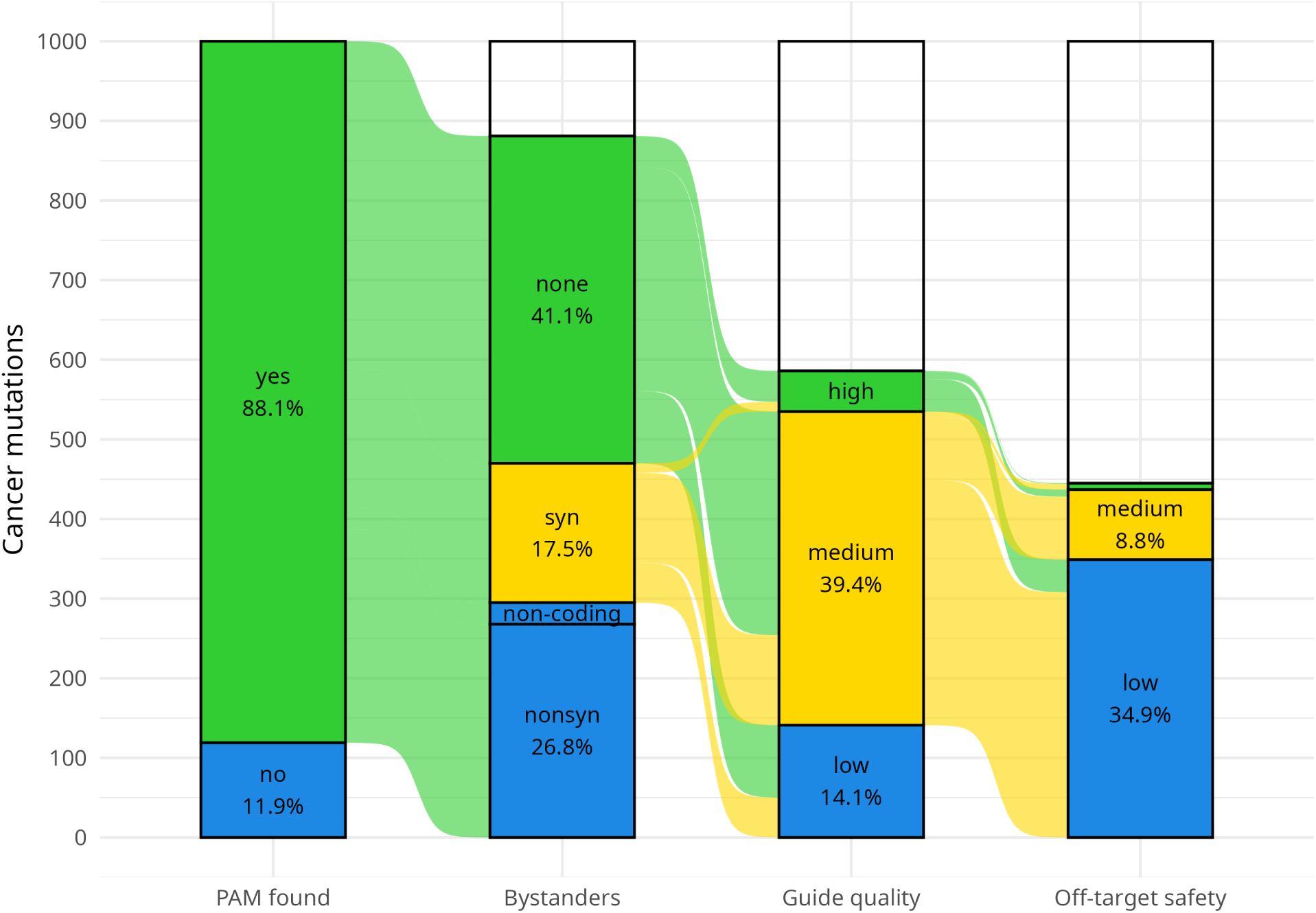
Relaxed-specificity NGN-PAM SpCas9 ABE attrition funnel for the top-1,000 ABE-editable cBioPortal mutations.

**Supplementary Figure S4:**
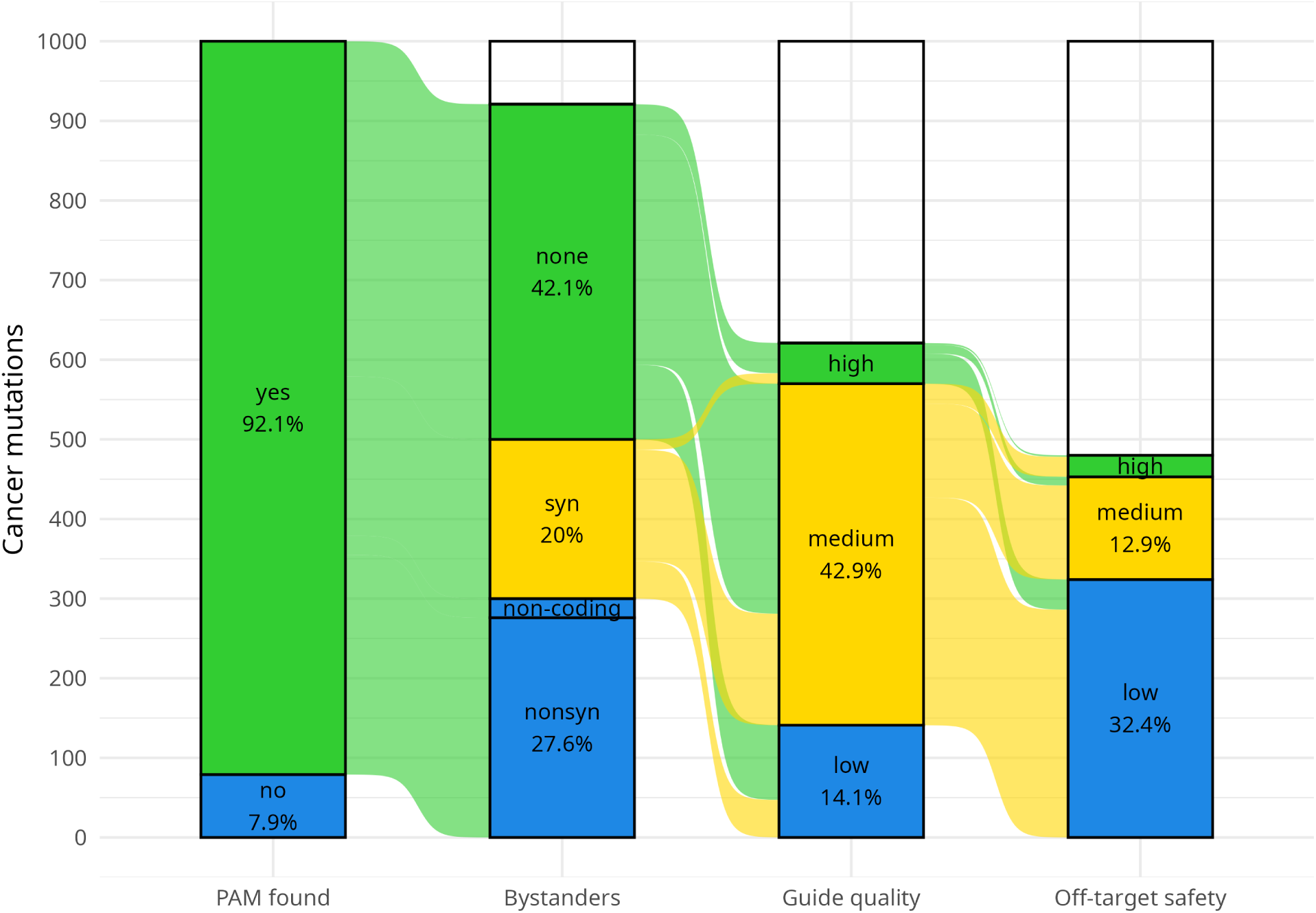
SaCas9 + SpCas9 + SpCas9-NG/SpG ABE (no SpRY) attrition funnel for the top-1,000 ABE-editable cBioPortal mutations.

**Supplementary Figure S5:**
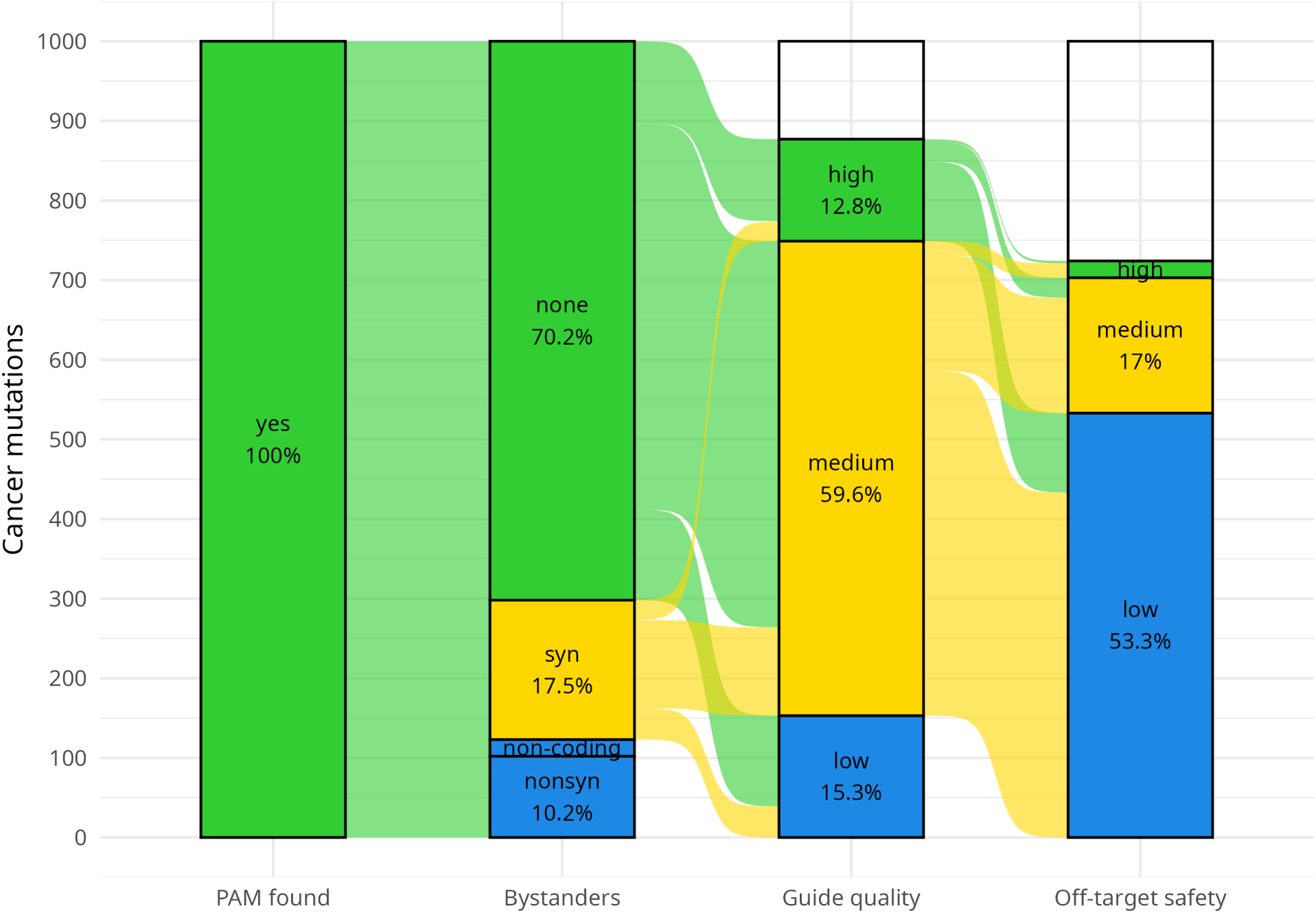
SpRY (NNN PAM) ABE attrition funnel for the top-1,000 ABE-editable cBioPortal mutations.

**Supplementary Figure S6:**
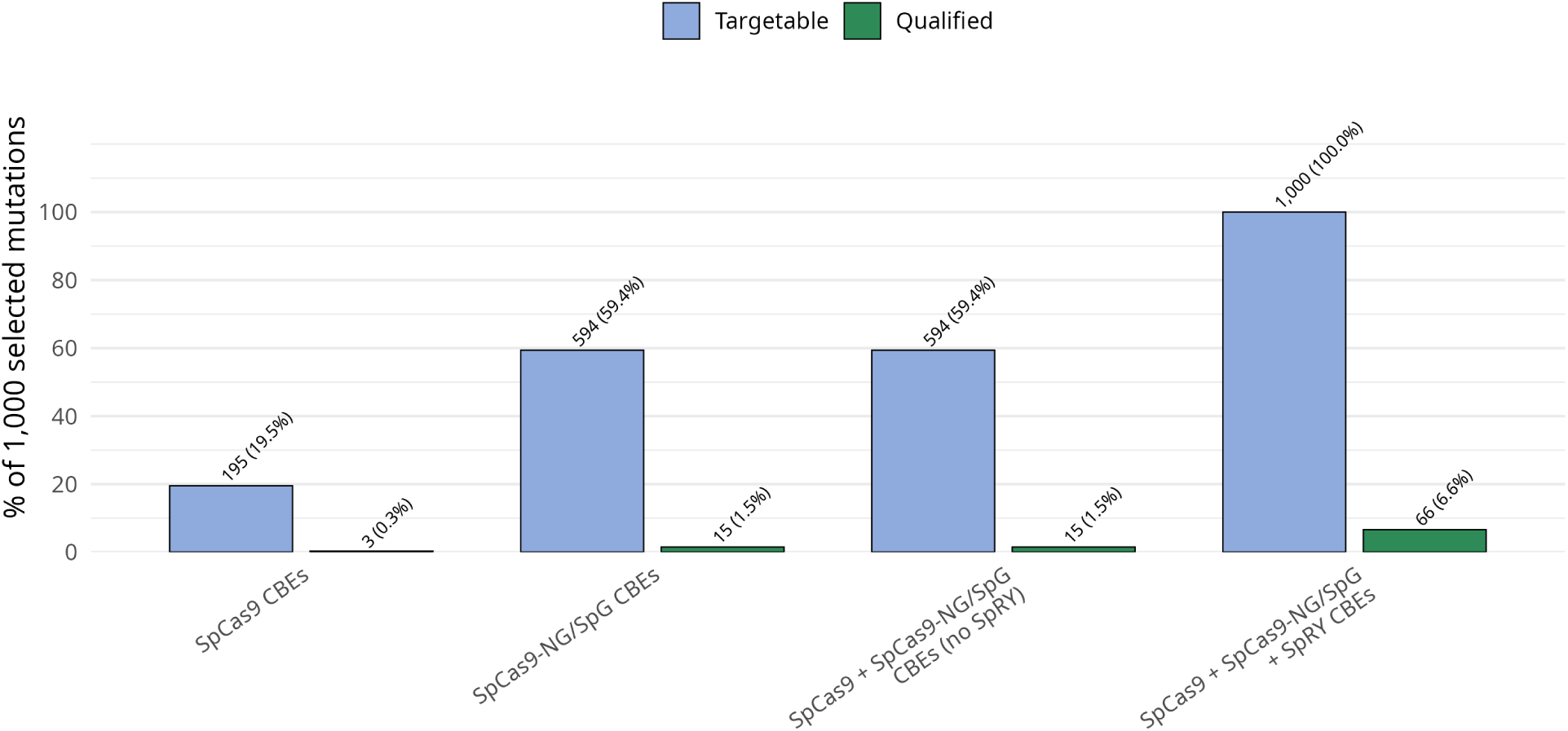
Targetability of the top-1,000 most frequent CBE-editable cBioPortal mutations across editor panels. Bars show the percentage of the 1,000 selected mutations that are targetable and the percentage of targets with at least one qualified guide, as defined for Figure 8. Labels above the bars give target counts and percentages.

**Supplementary Figure S7:**
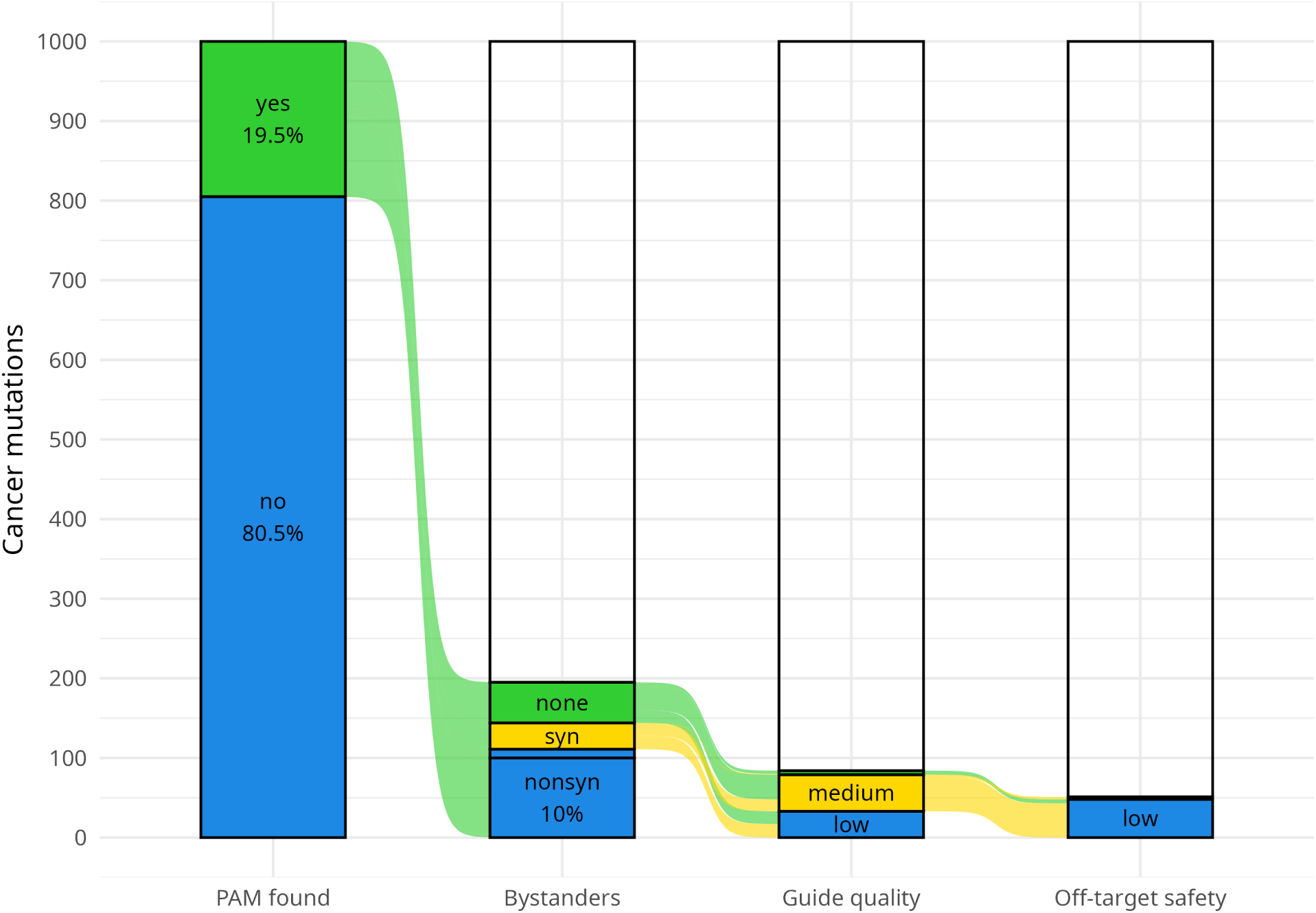
Attrition across filtering stages for the SpCas9 CBE panel (TadCBEd, CBE6d, TCBE-Umax, and TCBE-Umax-rest1). The total entering the funnel is 1,000 selected target mutations. Colors indicate favorable (green), intermediate (yellow), or unfavorable (blue) outcomes.

**Supplementary Figure S8:**
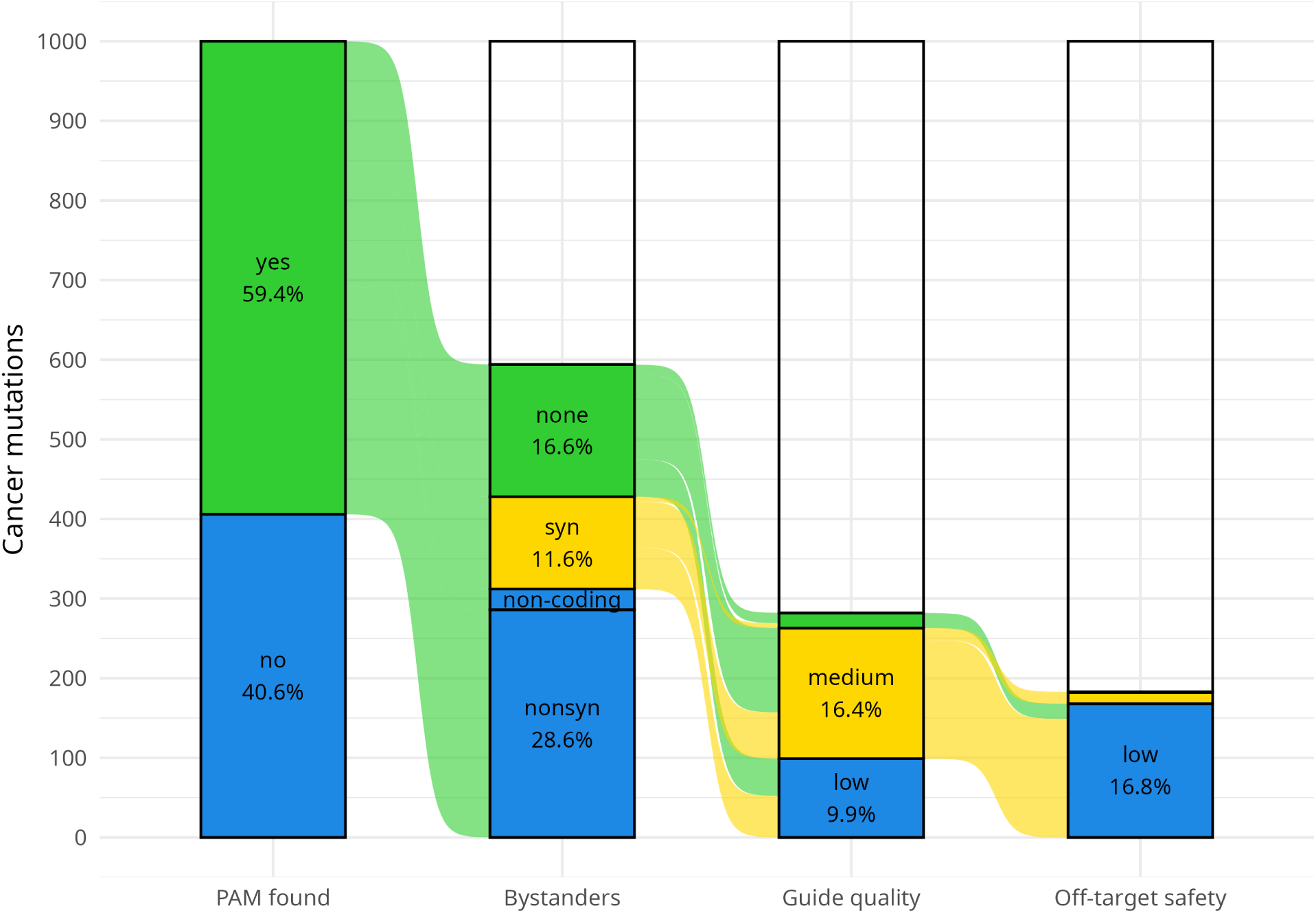
Relaxed-specificity NGN-PAM SpCas9 CBE attrition funnel for the top-1,000 CBE-editable cBioPortal mutations.

**Supplementary Figure S9:**
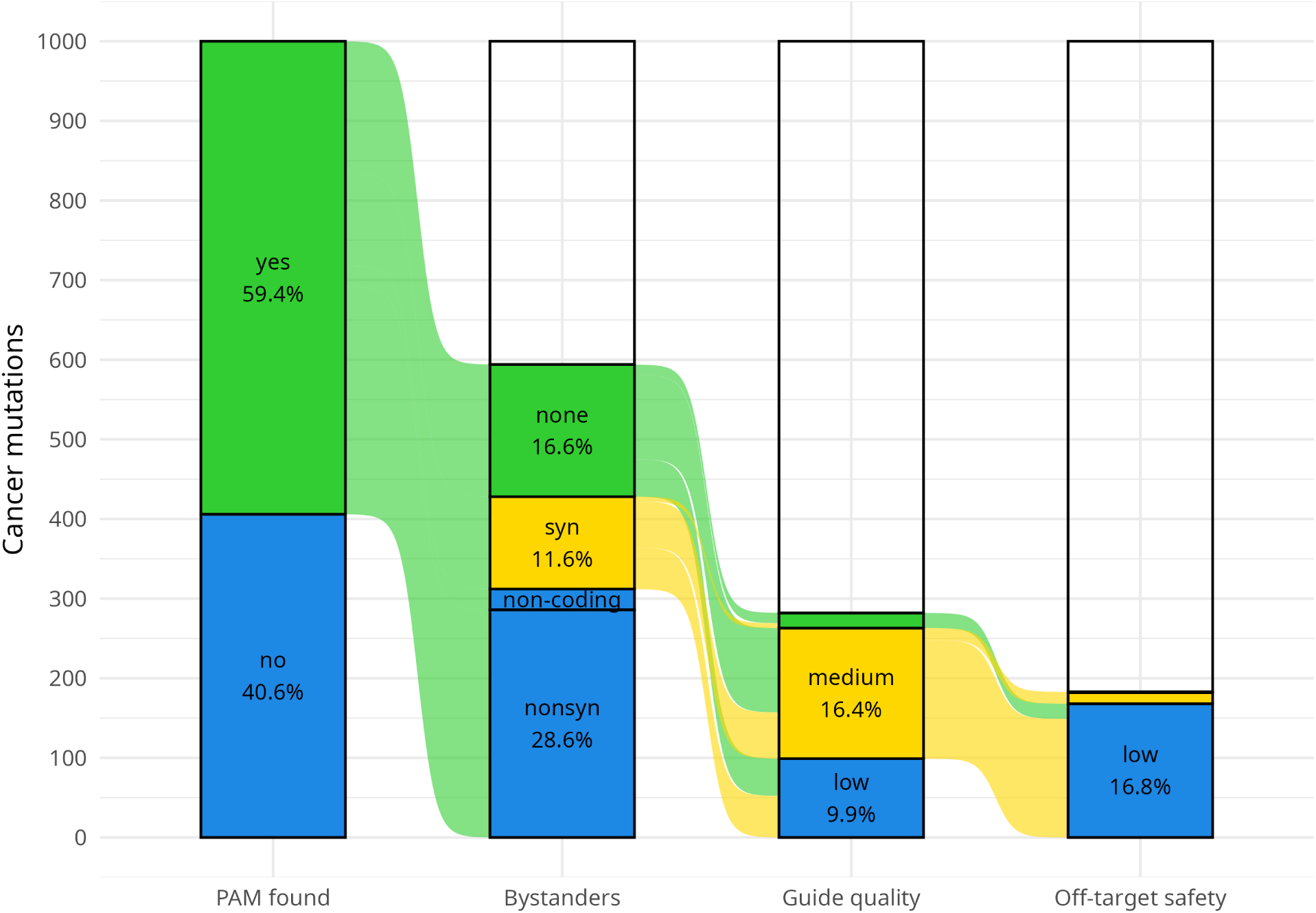
SpCas9 + SpCas9-NG/SpG CBE (no SpRY) attrition funnel for the top-1,000 CBE-editable cBioPortal mutations.

**Supplementary Figure S10:**
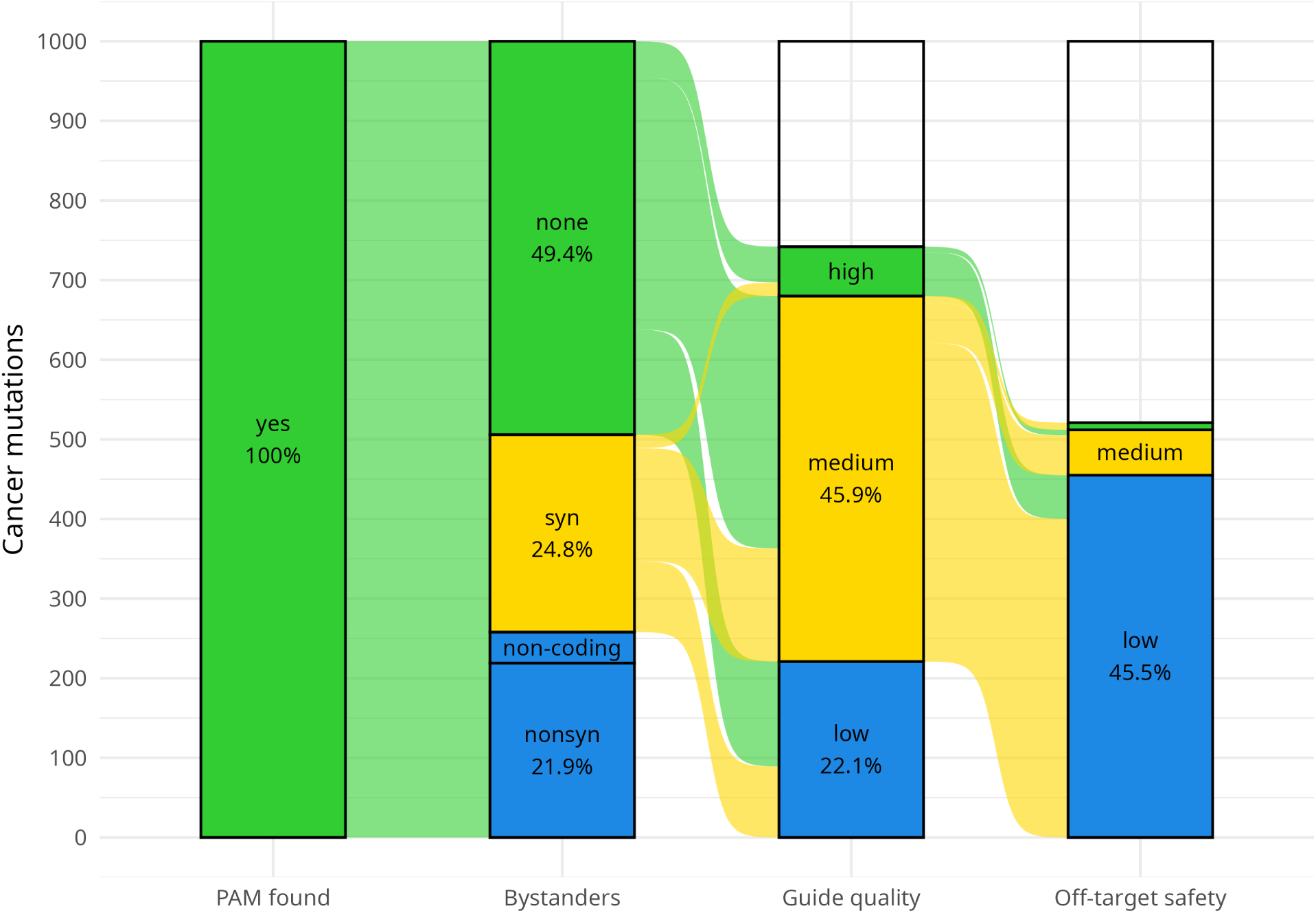
SpRY (NNN PAM) CBE attrition funnel for the top-1,000 CBE-editable cBioPortal mutations.

**Supplementary Figure S11:**
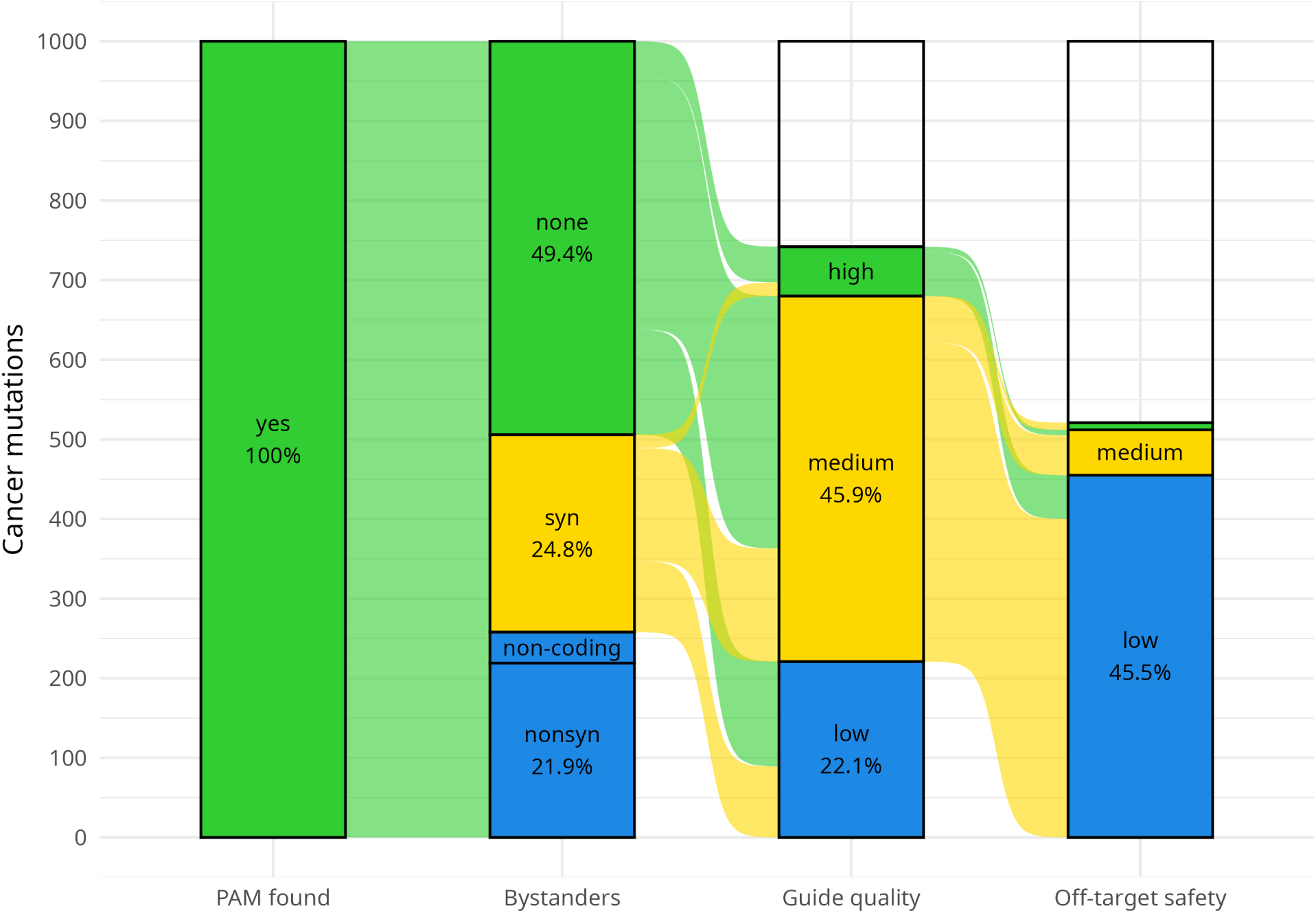
SpCas9 + SpCas9-NG/SpG + SpRY CBE attrition funnel for the top-1,000 CBE-editable cBioPortal mutations.

